# Trojan horse virus delivering CRISPR-AsCas12f1 controls plant bacterial wilt caused by *Ralstonia solanacearum*

**DOI:** 10.1101/2023.11.29.569319

**Authors:** Shiwen Peng, Yanan Xu, Hao Qu, Fushang Nong, Fangling Shu, Gaoqing Yuan, Lifang Ruan, Dehong Zheng

## Abstract

Plant bacterial wilt caused by the *Ralstonia solanacearum* species complex results in huge food and economic losses. Accordingly, the development of an effective control method for this disease is urgently required. Traditional lytic phage biocontrol methods have inherent limitations. However, filamentous phages, which do not lyse host bacteria and exert minimal burden, offer a potential solution. A filamentous phage RSCq that infects *R. solanacearum* was isolated in this study through genome mining. We constructed engineered filamentous phages based on RSCq by employing our proposed approach with a wide applicability to non-model phages, enabling the infection of *R. solanacearum* in medium and soil and delivering exogenous genes into bacterial cells. Similar to the Greek soldiers hidden within the Trojan horse, CRISPR-AsCas12f1 gene editing system that targets the key virulence regulator gene *hrpB* was implanted into the engineered phage, generating the engineered phage RSCqCRISPR-Cas. Our findings demonstrated that RSCqCRISPR-Cas could disarm the key “weapon”, *hrpB*, of *R. solanacearum*, in medium and in plants. Remarkably, pretreatment with RSCqCRISPR-Cas significantly controlled tobacco bacterial wilt, highlighting the potential of engineered filamentous phages as promising biocontrol agents against plant bacterial wilt and other bacterial diseases.

## Introduction

An estimated 821 million people–approximately 1 out of every 9 people in the world are undernourished ^1^. Moreover, the world’s population is expected to reach 9.6 billion by 2050, and additional global food supply will be needed to meet the increasing demands ^2^. However, plant diseases cause around 10%–20% of global food production loss every year ^3, 4^. For instance, the plant bacterial pathogen *Ralstonia solanacearum* species complex infects more than 400 plant species from over 50 families, and losses due to the disease on potatoes alone were estimated at US$1 billion each year worldwide ^5^. Nevertheless, efficient control methods for managing plant bacterial wilt caused by the *R. solanacearum* species complex remain highly limited.

Efficient control methods for plant diseases are of utmost importance to ensure optimal crop production and meet the ever-increasing demands for food. Among the different strategies, the utilization of phages, which are viruses that infect bacteria, exhibits considerable promise. Phage therapy, an environment-friendly control strategy, has been frequently reported in plant bacterial disease control ^6^. For example, phage ΦPD10.3 and ΦPD23.1 reduced the severity of potato soft rot caused by *Pectobacterium carotovorum* by 80%-95% ^7^. In another study, a phage cocktail that consisted of six phages effectively suppressed symptom development of leek bacterial blight caused by *Pseudomonas syringae* ^8^. Similarly, increasing the number of *R. solanacearum* phages in various combinations decreased the incidence of tomato bacterial wilt disease by up to 80% ^9^. However, limitations still impede the widespread application of phage therapy in crop production. The host specificity of phages is a major disadvantage that may be partially overcome by the development of phage cocktails. Moreover, the emergence of bacterial resistance to phage infection poses challenges to the continuous use of phage treatments. In addition, phage sensitivity to ultraviolet (UV) light and certain soil conditions cause phage decline after application, undermining the biocontrol effect ^6^.

In general, strictly lytic phages are preferred for biocontrol applications. However, temperate phages, which are highly abundant, should not be overlooked ^10^. Filamentous phages belong to the *Inoviridae* family of phages with small single-stranded DNA (ssDNA) genomes packaged within filament-like virions. In contrast with lytic phages, filamentous phages do not lyse or otherwise kill the host bacterium, but instead, egress from the host cell, imposing minimal burden on bacteria ^11^. *R. solanacearum* loses virulence on tomato plants under the infection of filamentous phages RSM1 and RSM3^12^. Although many other filamentous phages have no biocontrol effect and even enhance the virulence of phytopathogenic bacteria ^13^, the huge diversity of filamentous prophages integrated into bacterial genome and the small genome size make filamentous phages promising genetically engineered biocontrol reagents and biotechnological tools ^10, 14^. Moreover, bacteria infected by filamentous phages continue to produce infectious phage particles, which can potentially counteract the influence of UV light and other hostile environmental conditions. Consequently, filamentous phages offer a potential solution to overcome the limitations of lytic phage biocontrol.

In the mythological tale of the Trojan horse, Greek soldiers concealed themselves inside a large wooden horse to gain entry into the city of Troy and secure victory in the Trojan War, our study employed a similar strategy. We constructed filamentous phage-based “Trojan horses”, metaphorical “gifts”, aimed at the pathogenic bacteria *R. solanacearum*. Drawing inspiration from the legendary soldiers hidden inside the Trojan horse, we utilized the clustered regularly interspaced short palindromic repeat and CRISPR-associated proteins (CRISPR-Cas) system, which specifically targets the key virulence regulator gene *hrpB* of *R. solanacearum*. The CRISPR-Cas system was successfully delivered through the engineered phages into *R. solanacearum* cells, leading to efficient control of plant bacterial wilt caused by this pathogen. In essence, our study harnessed the concept of the Trojan horse to combat this disease effectively.

## Results

### Prophage mining in the genomes of *R. solanacearum*

Many filamentous phages can integrate into the host chromosome and replicate with the bacterial genome ^11^. In this study, 50 phylotype I, 9 phylotype II, 3 phylotype III, and 12 phylotype IV *R. solanacearum* strains with publicly available completed genomes were analyzed to evaluate the diversity of integrated prophages by using the phage search tool PHASTER ^15^. As shown in Figure 1a and Table S1, at least 1 intact prophage sequence was found in 63 of the 74 *R. solanacearum* strains, resulting in 152 intact filamentous prophage sequences in the total strains. Among these 152 sequences, 50 encode filamentous phages, on the basis of the result of the “Most Common Phage”, which is an important term in PHASTER defined by the phage(s) with the highest number of proteins most similar to those in the identified prophage. Filamentous phage RSS0 (sequence accession: NC_019548) that infects *R. solanacearum* was most frequently identified as the “Most Common Phage” (29/50), followed by RSM3 (7/50) and PE226 (6/50) that infects *R. solanacearum* (Figure 1b and Table S1). The 50 filamentous prophage sequences are distributed in 42 *R. solanacearum* strains (36 phylotype I strains, 4 phylotype II strains, and 2 phylotype III strains). These results suggest that filamentous prophages are distributed widely throughout *R. solanacearum* phylotypes I, II, and III strains, underscoring the feasibility of isolating filamentous phages that infect *R. solanacearum* through genome mining approaches.

**Figure 1.**
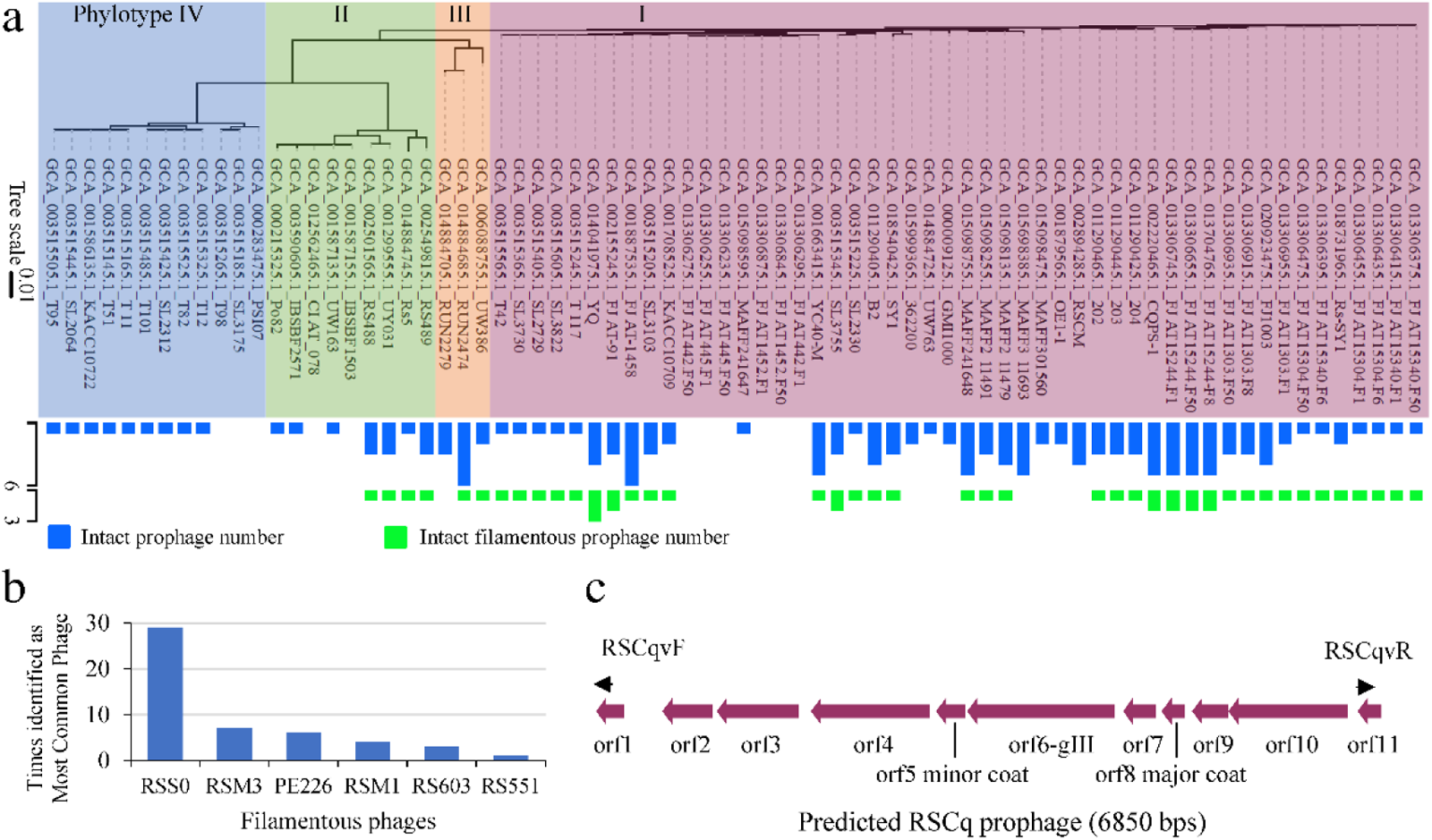
Prophage mining in the genomes of *R. solanacearum*. **a**. Prophages was predicted via PHASTER. The genomic phylogenetic tree of *R. solanacearum* strains was constructed via GToTree based on a single-copy gene set. The number of predicted prophages and the phylogenetic tree were visualized via tvBOT. **b**. Times of filamentous phages identified as “Most Common Phage” which is defined by the phage(s) with the highest number of proteins most similar to those in the identified prophage. **c**. A filamentous prophage sequence was identified in the *R. solanacearum* phylotype I strain Cq05 isolated from grafted chiehqua at Nanning, Guangxi Province, China.

*R. solanacearum* phylotype I (*R. pseudosolanacearum*) strain Cq05 isolated from grafted *chiehqua* in Nanning, Guangxi Province, China, was genome-sequenced to discover filamentous phage and the genome sequence was deposited at GenBank (BioProject ID PRJNA974909). As indicated in Table 1, five prophage regions, with three intact regions (region1, region2, and region5), were identified in the genome of Cq05. The “Most Common Phage” of region1 and region2 are RSY1 and RSA1, respectively. RSY1 and RSA1 belong to the *Myoviridae* family of phages that infect *R. solanacearum*. As shown in Figure 1c, the 6.8kb region5 sequence in contig JASKHZ010000087.1 was predicted to be an intact prophage. This prophage sequence encodes 11 open reading frames (ORFs), all of which are homologous to that of the reported *R. solanacearum* filamentous phage RSS1 ^16^ (sequence accession: NC_008575). *orf5* encodes a putative minor coat protein, while *orf8* encodes a putative major coat protein (pVIII) of filamentous phages. *orf6* encodes a putative pIII protein, which together with pVI, caps the terminal end of filamentous phages. These findings suggest that prophage region5 may encode a potential filamentous phage, which is named as RSCq.

**Table 1.**
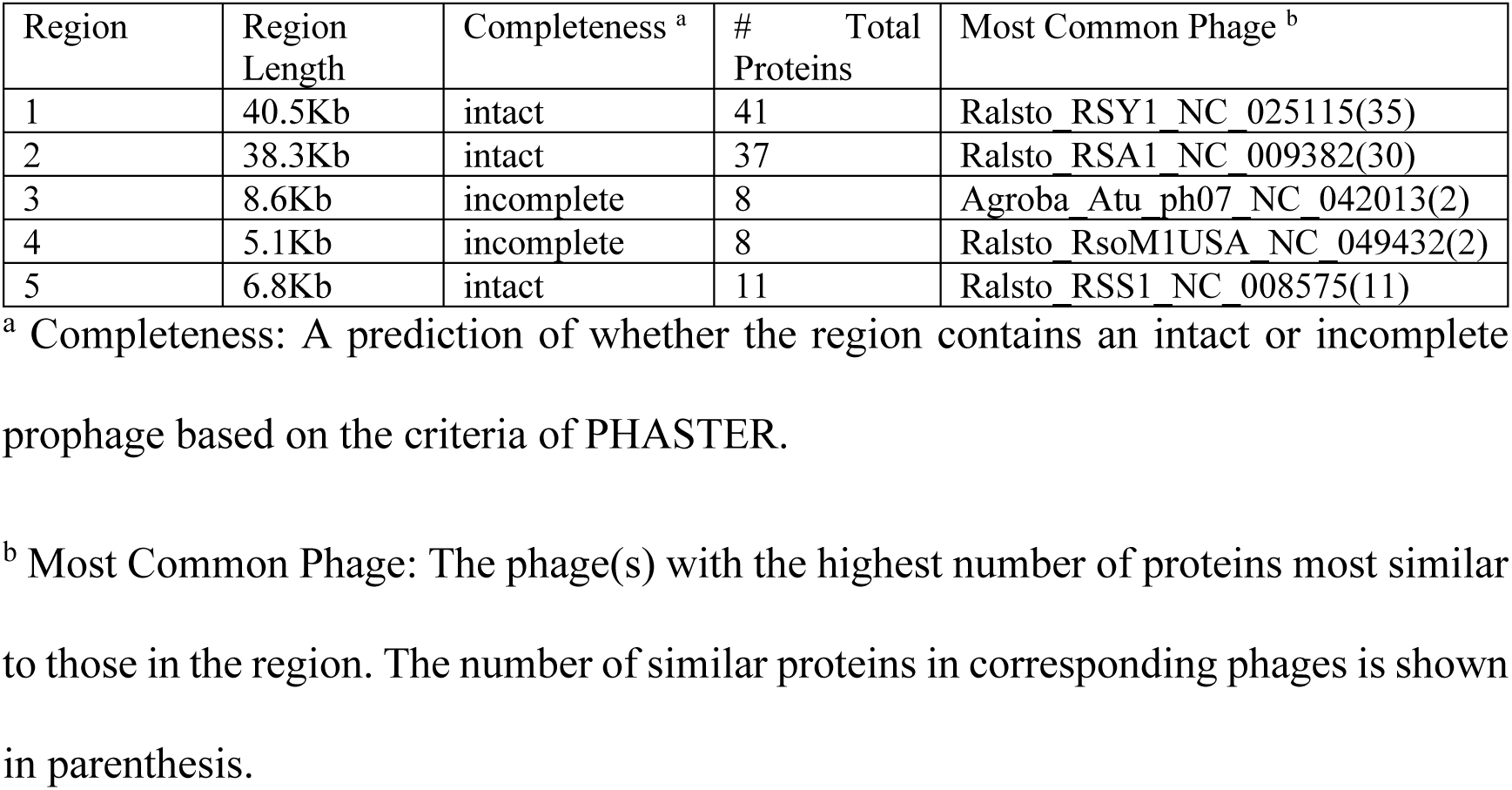
Prophage regions identified in *R. solanacearum* Cq05.

| Region | Region Length | Completeness <sup>a</sup> | # Total Proteins | Most Common Phage <sup>b</sup> |
| --- | --- | --- | --- | --- |
| 1 | 40.5Kb | intact | 41 | Ralsto_RSY1_NC_025115(35) |
| 2 | 38.3Kb | intact | 37 | Ralsto_RSA1_NC_009382(30) |
| 3 | 8.6Kb | incomplete | 8 | Agroba_Atu_ph07_NC_042013(2) |
| 4 | 5.1Kb | incomplete | 8 | Ralsto_RsoM1USA_NC_049432(2) |
| 5 | 6.8Kb | intact | 11 | Ralsto_RSS1_NC_008575(11) |

<sup>a</sup> Completeness: A prediction of whether the region contains an intact or incomplete prophage based on the criteria of PHASTER.
<sup>b</sup> Most Common Phage: The phage(s) with the highest number of proteins most similar to those in the region. The number of similar proteins in corresponding phages is shown in parenthesis.

### Isolation and characterization of the filamentous phage RSCq

To isolate the filamentous phage RSCq, we took advantage of the cooperative relationship between filamentous phages and their bacterial hosts, in which phages are continuously secreted from the hosts. We initially detected the presence of RSCq in the culture supernatant of Cq05 via double agar overlay plaque assay using *R. solanacearum* GMI1000 as host bacteria. Small and turbid plaques were observed, as illustrated in Figure 2a. Subsequently, two rounds of single plaque picking up and infection were performed to purify plaque. The isolated filamentous phage RSCq was cultured using GMI1000 as host bacteria. RSCq in GMI1000 was verified via polymerase chain reaction (PCR) by using primers RSCqvF and RSCqvR (Table S3). In Figure 2b, the filamentous phage RSCq was inoculated with tested host bacteria at a multiplicity of infection (MOI) of 10. *R. solanacearum* phylotype I Bg06 and Bg07 are isolated from bitter gourd in Guangxi Province, China. The presence of RSCq significantly delayed the growth of *R. solanacearum* GMI1000, Bg06, and Bg07, but did not lyse them. This finding is consistent with the biological features of filamentous phages.

**Figure 2.**
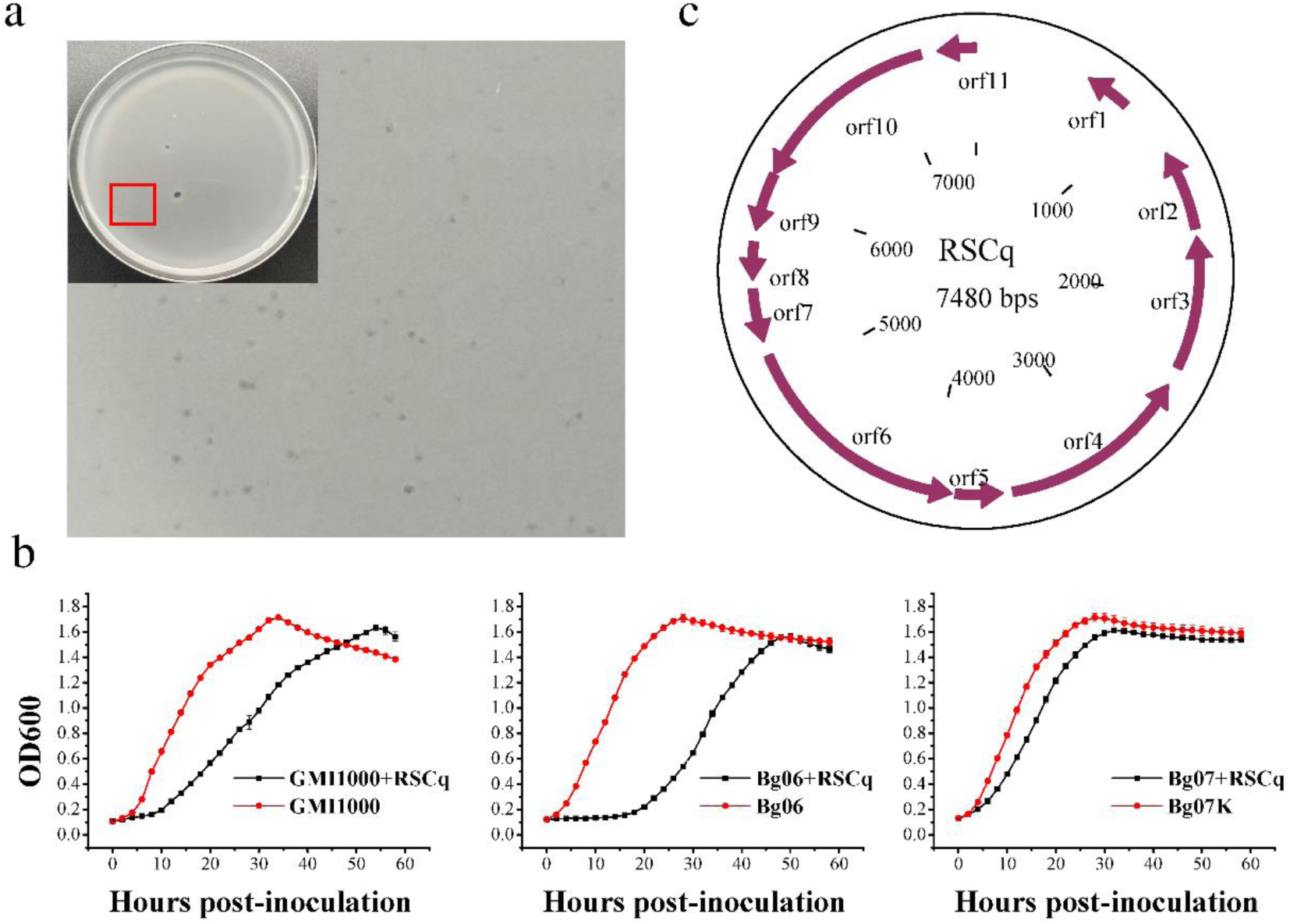
Isolation and characterization of the filamentous phage RSCq. **a**. Double agar overlay plaque assay of *R. solanacearum* Cq05 culture supernatant by using *R. solanacearum* GMI1000 as host bacteria. **b**. Effect of filamentous phage RSCq infection on the growth of *R. solanacearum* GMI1000, Bg06, and Bg07. The growth curves of *R. solanacearum* strains with or without RSCq infection were measured by monitoring bacterial growth (A600) via Bioscreen C Pro. The error bar is represented by the standard deviation of three technical repeats. The growth curves were assayed three times independently **c**. The replicative form genome of filamentous phage Cq05.

Filamentous phages typically have a circular ssDNA genome within their virions, but a double-stranded replicative form (RF) of the genome exists in the host bacteria during the phage life cycle ^11^. The RF DNA of RSCq was then extracted via the plasmid DNA purification procedure. To verify the RF DNA of RSCq and compare the genome sequence of RSCq with the predicted genome, we amplified and sequenced the flanking sequence of the junction site by using primers that bind to *orf1* and *orf11* (Figure 1c). Unexpectedly, the amplification product is 925 bps in length, which is larger than the deduced 295 bps based on the PHASTER analysis. This discrepancy suggested that the genome of RSCq is larger than initially predicted. The genome was then corrected to 7480 bps on the basis of the sequence of this 925 bps DNA (Figure 2c), and confirmed via whole genome Sanger sequencing (10.6084/m9.figshare.24473443). The genome sequence of the filamentous phage RSCq was deposited in GenBank (Accession number: OR088903).

### Construction of engineered phages based on RSCq

We assayed the effect of phage RSCq infection on the virulence of *R. solanacearum* through the stem injection of tomato and tobacco plants. No remarkable virulence difference was observed between phage-infected and uninfected *R. solanacearum* (Figure 7 and Supplementary Figure 4). When *R. solanacearum* infected tobacco via natural inoculation, RSCq infection delayed bacterial wilt symptoms but resulted in a similar final disease index of tobacco plants (Figure 8). This result prompted us to construct engineered phages that are capable of delivering biocontrol factors.

We proposed a novel method for constructing engineered filamentous phages that involved propagating RSCq RF DNA as an independently replicating plasmid in *Escherichia coli* by using Tn5 transposase. As depicted in Figure 3a, the transposon of the EZ-Tn5™ <R6Kγori/KAN-2> Insertion Kit (Lucigen, Wisconsin, USA) was modified. The modified transposon contained the *eYFP* gene controlled by the lac promoter, which was inserted between the transposase recognition sequence (ME) and the kanamycin resistance gene (KanR). The modified transposon was then randomly inserted into RSCq RF DNA in vitro by Tn5 transposase. The resulting transposon-inserted plasmid library was electrotransformed into *R. solanacearum* GMI1000, followed by engineered phage screening on the basis of the growth inhibition effect. The supernatant of three transformants showed a growth inhibition effect on GMI1000, based on the result of the inhibition zone (Figure 3b) and the growth curve (Figure 3c). This finding suggests that the engineered phages were secreted from these transformants and named RSCqYFP01, RSCqYFP02, and RSCqYFP03.

**Figure 3.**
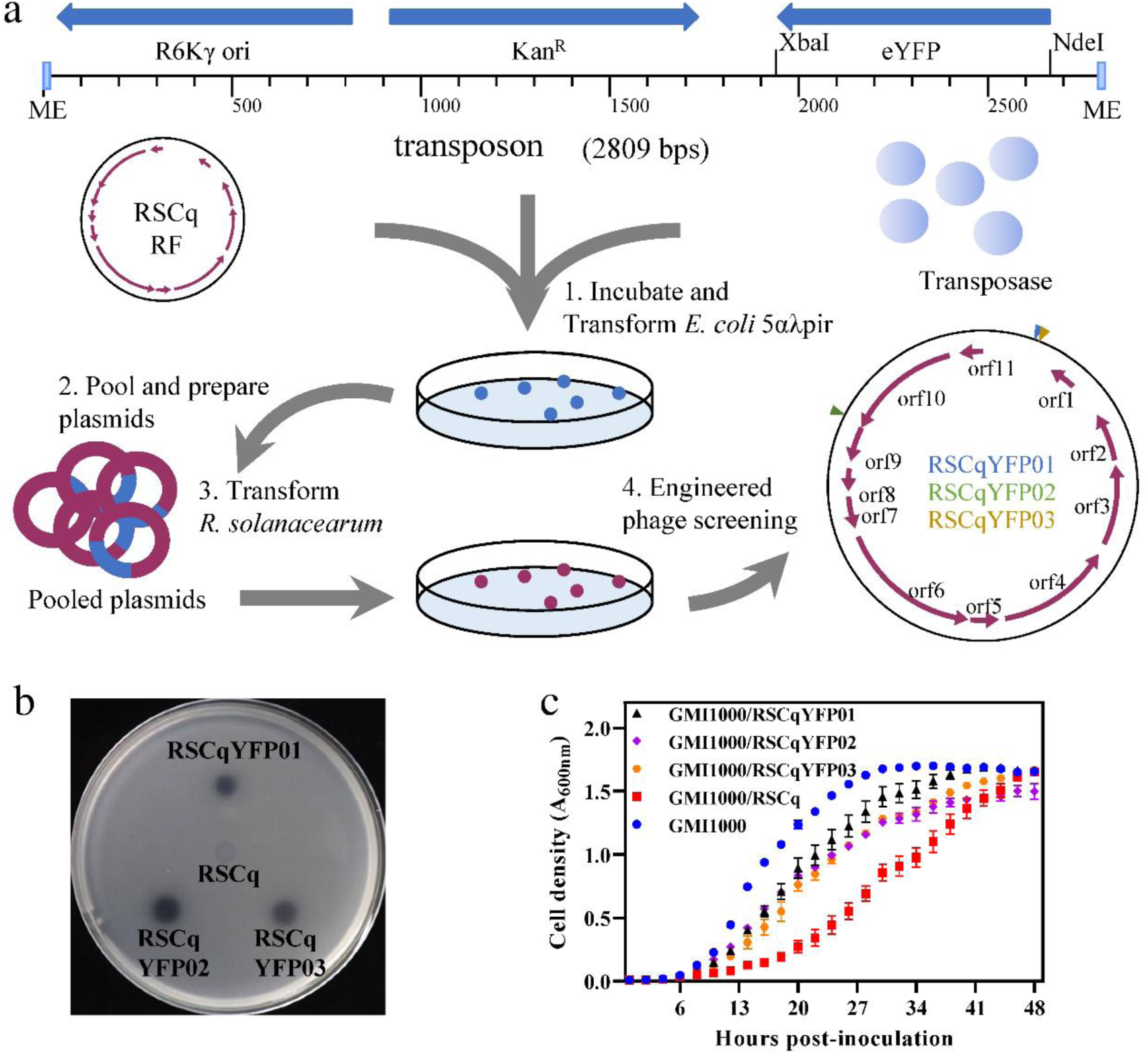
Construction of engineered phages based on RSCq a. Construction procedure of engineered phages based on RSCq. The modified transposon containing an *eYFP* gene was randomly inserted into RSCq replicative form DNA in vitro by Tn5 transposase, generating a plasmid library. The plasmids transposon inserted library was then transformed into *R. solanacearum* GMI1000 to recover engineered phages. **b.** Inhibition zone of engineered phages on *R. solanacearum* GMI1000. **c**. Growth curve of *R. solanacearum* GMI1000 infected by engineered phages. The error bar is represented by the standard deviation of three technical repeats. The growth curves were assayed three times independently

The inserted site in the genome of the engineered phages was determined via Sanger sequencing by using primers that bind on the transposon. Consequently, the insertion sites were identified for each engineered phage. In the case of RSCqYFP01, the transposon was inserted into the noncoding regions downstream of *orf1*, particularly at positions 553-554 bp of the RSCq RF DNA. For RSCqYFP03, the transposon was inserted at positions 561-562 bp. RSCqYFP02 had its transposon inserted at positions 6168-6169 bp of the RSCq RF DNA, which is located at the 3’ end of *orf10*.

### Infectious feature of engineered phage RSCqYFP01

Infectious capability was assayed to confirm the engineered filamentous phages. The engineered filamentous phages RSCqYFP01, RSCqYFP02, and RSCqYFP03 were inoculated with *R. solanacearum* GMI1000 in BG medium. The kanamycin resistance gene present in the engineered filamentous phages enabled the identification of infected *R. solanacearum* cells. As depicted in Figure 4a, bacterial culture at various time points (0, 4, 8, and 12 h) after inoculation was streaked onto BG agar medium with or without kanamycin, and the result showed that *R. solanacearum* acquired kanamycin resistance when co-cultured with RSCqYFP01, RSCqYFP02, or RSCqYFP03. The infectious efficiency 12 h post-inoculation was quantified by the colony-forming unit on BG agar medium with or without kanamycin. As shown in Figure 4b, more than 85% of *R. solanacearum* cells exhibited resistance to kanamycin after 12 h of engineered filamentous phage infection. These findings signify the efficient infectivity of the engineered filamentous phages in host bacteria. Among the engineered phages, RSCqYFP01 was selected for further research. Supplementary Figure 1 provides the map of RSCqYFP01, while additional details regarding the sequence can be found in Supplementary Note 1.

**Figure 4.**
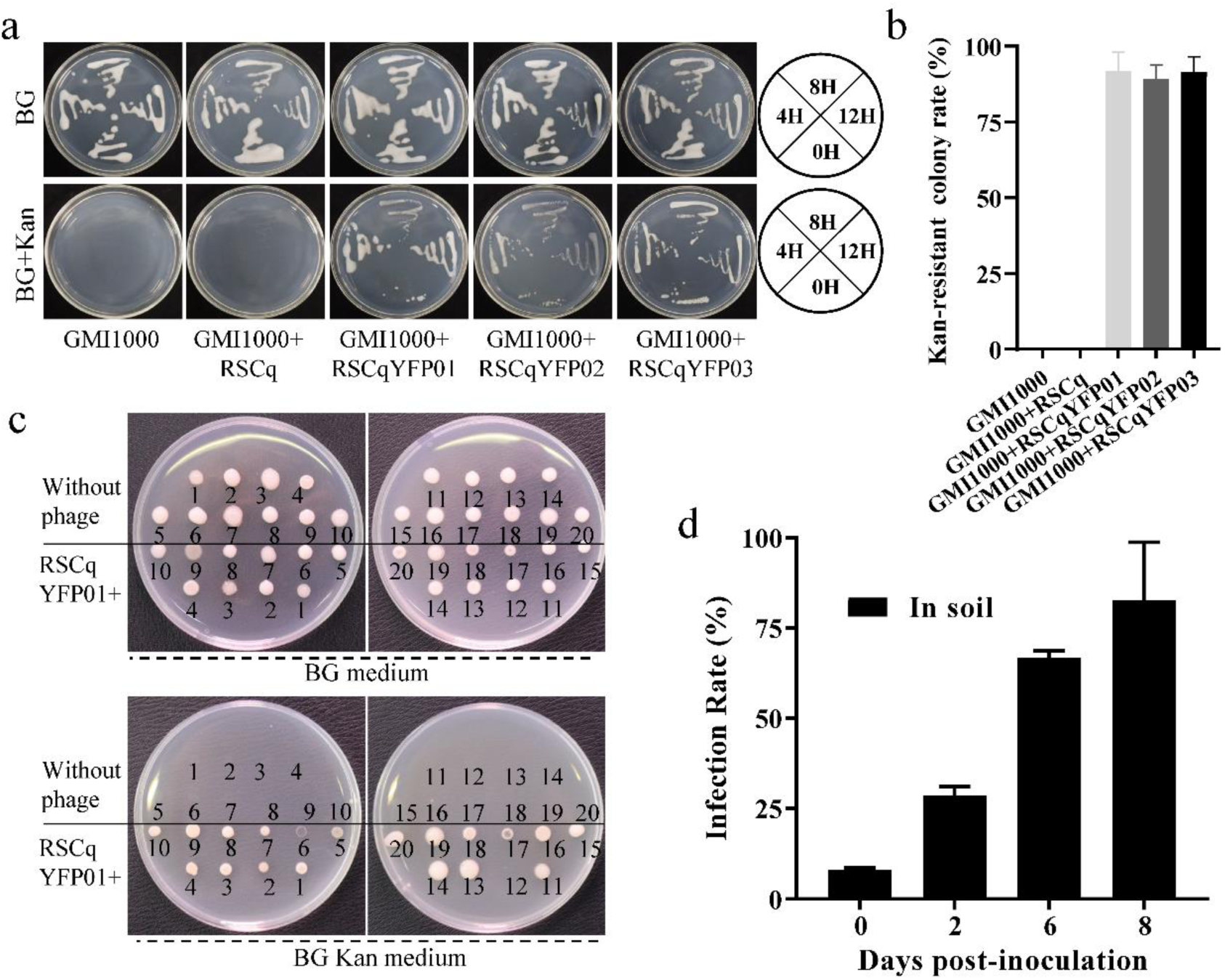
Infectious feature of engineered phage. **a**. *R. solanacearum* streaked on BG agar medium with or without kanamycin at 0, 4, 8, and 12 h after the infection of the phage RSCq or the engineered phages RSCqYFP01, RSCqYFP02, or RSCqYFP03. **b.** The kanamycin-resistant colony rate of *R. solanacearum* plating on BG agar medium with or without kanamycin 12 h post-infection of the phage RSCq or the engineered phages. The error bar is represented by the standard deviation of three technical repeats. This experiment was performed three times independently. **c**. *R. solanacearum* on BG agar medium with or without kanamycin at 12 h after co-culturing the engineered phage RSCqYFP01 and the tested *R. solanacearum* strains (Supplementary Table 2). **d**. The infection rate of RSCqYFP01 on *R. solanacearum* in soil substrate. The error bar is represented by the standard deviation of three technical repeats. This experiment was performed three times independently.

In addition to *R. solanacearum* GMI1000, 19 *R. solanacearum* phylotype I strains isolated from diverse plant hosts in various locations within Guangxi, China, were tested to determine the host range of the engineered filamentous phage. Detailed information regarding the 19 strains is provided in Supplementary Table 2. RSCqYFP01 and *R. solanacearum* strains were cocultured in BG medium for 12 h, followed by kanamycin resistance assay. As shown in Figure 4d, RSCqYFP01 successfully infected 19 out of 20 of the tested strains, indicating that the engineered filamentous phage exhibits a broad host range on *R. solanacearum* phylotype I strains.

Furthermore, the infection efficiency of the engineered filamentous phage in soil was also assayed to evaluate its potential for biocontrol applications. *R. solanacearum* GMI1000 was introduced into a soil substrate at a final concentration of 10^8^ colony-forming units per gram of substrate, followed by RSCqYFP01 infection after 1 day of *R. solanacearum* watering. As shown in Figure 4d, 82.6% of *R. solanacearum* cells in the soil were infected after 8 days of RSCqYFP01 treatment. The “Trojan horse” can be efficiently implanted into *R. solanacearum* in a medium and in soil environment.

### Target gene can be delivered to host bacteria by engineered phage

As mentioned above, an *eYFP* gene was introduced into the transposon when constructing the engineered phages. The inoculation of RSCqYFP01 and *R. solanacearum* GMI1000 in BG medium for 48 h resulted in a yellow-green fluorescence signal in the infected bacteria under a fluorescence microscope (Figure 5a). By contrast, no fluorescence signal was observed in *R. solanacearum* infected by the parent phage RSCq. Measurements of fluorescence density using a detection reader showed only background fluorescence in non-infected and RSCq-infected strains; meanwhile, high fluorescence density was detected in the RSCqYFP01-infected strain (Figure 5b).

**Figure 5.**
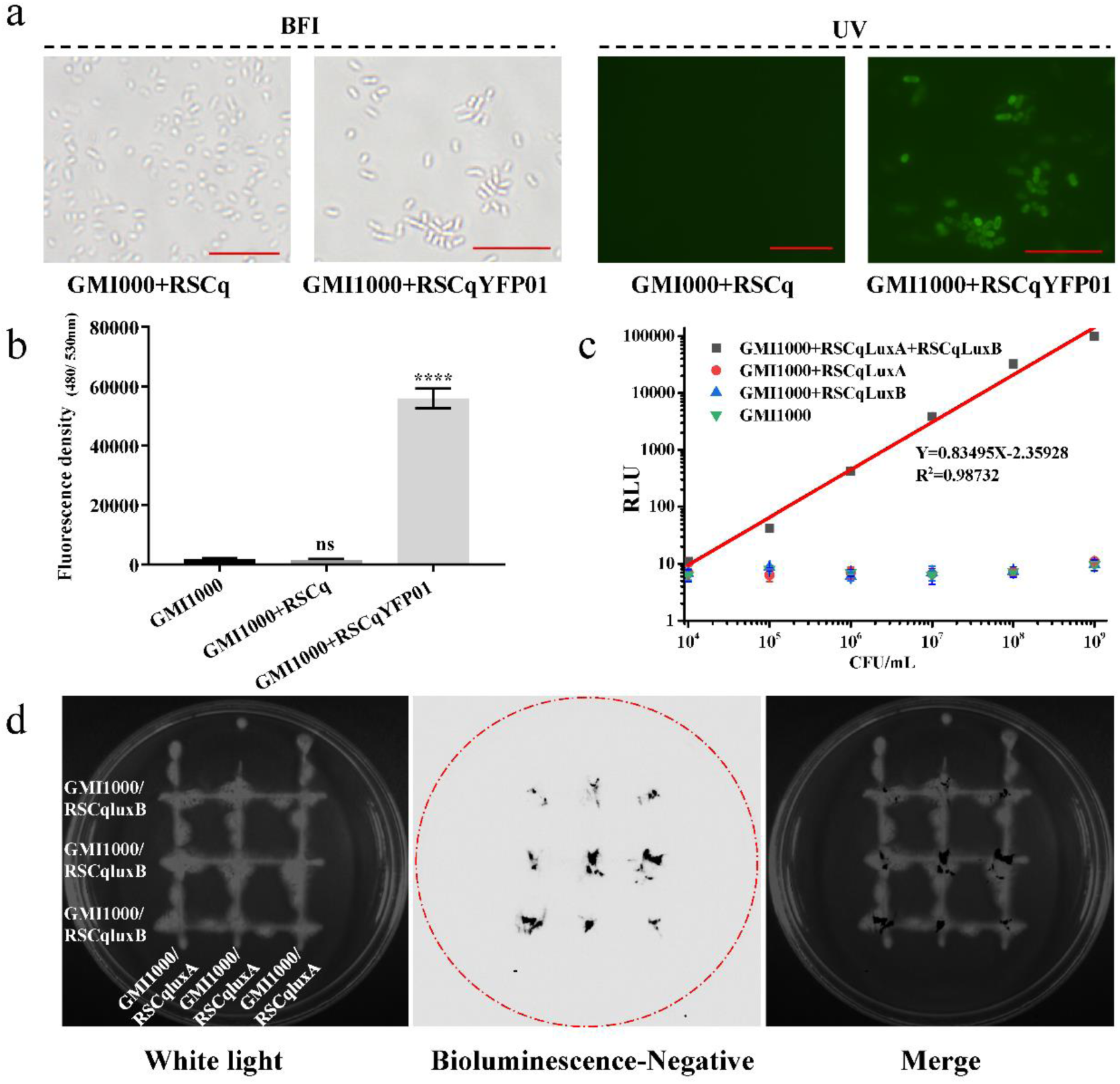
Target genes can be delivered into host bacteria by engineered phage. **a**. Engineered filamentous phage RSCqYFP01 infected *R. solanacearum* GMI1000 under a fluorescence microscope. BFI, bright field images. UV, images under ultraviolet excitation. **b**. The fluorescence density of the engineered filamentous phage RSCqYFP01 infected *R. solanacearum* GMI1000. The error bar is represented by the standard deviation of three technical repeats. This experiment was performed three times independently. **c**. Relative light units of *R. solanacearum* GMI1000 co-infected by engineered filamentous phages RSCqluxA and RSCqluxB. RSCqluxA or RSCqluxB infected *R. solanacearum* GMI1000, and GMI1000 without phage infection were used as controls. The error bar is represented by the standard deviation of three technical repeats. This experiment was performed three times independently. **d**. Luminescence imaging of the engineered filamentous phages RSCqluxA- and RSCqluxB-infected *R. solanacearum* (RSCqluxA/GMI1000 and RSCqluxB/GMI1000, respectively) on BG agar medium. Left panel, image of RSCqluxA/GMI1000 and RSCqluxB/GMI1000 under white light. Middle panel, the negative image of bioluminescence in the dark. Right panel, merged image.

We replaced the *eYFP* gene with *luxA* and *luxB* genes, generating engineered phage RSCqluxA and RSCqluxB, respectively, to confirm the delivery and expression of the exogenous gene further. The LuxAB heterodimeric enzyme, but not LuxA or LuxB alone, catalyzes the bioluminescence reactions emitting blue-green light ^17^. *R. solanacearum* GMI1000 was co-infected by engineered phages RSCqluxA and RSCqluxB in BG liquid medium for 24 h. The infected bacteria were subject to gradient dilution, and bioluminescence was measured after adding the luminescent substrate aldehyde. As shown in Figure 5c, an intense bioluminescent signal was detected in RSCqluxA/RSCqluxB co-infected GMI1000, but not in wild-type GMI1000, RSCqluxA- or RSCqluxB-infected GMI1000. The relative light units (RLUs) of RSCqluxA/RSCqluxB co-infected GMI1000 were correlated linearly with bacterial concentration. The infected bacteria were also diluted and plated on BG agar medium. The resulting colonies were imaged under a luminescence imaging system after spraying the luminescent substrate aldehyde. The luminescence of colonies indicated that RSCqluxA and RSCqluxB can co-infect a single *R. solanacearum* cell (Supplementary Figure 2).

RSCqluxA- and RSCqluxB- infected *R. solanacearum*, namely RSCqluxA/GMI1000 and RSCqluxB/GMI1000, respectively, were streaked on BG agar medium by crossing each other. Luminesce can be detected at the intersection of RSCqluxA/GMI1000 and RSCqluxB/GMI1000 after spraying of the luminescent substrate aldehyde (Figure 5d), suggesting engineered phages can be continuously secreted from the infected *R. solanacearum* cells and infect other surrounding host cells. This finding demonstrates that exogenous genes integrated with the engineered phage genome can be delivered and expressed efficiently in the host bacterial cells along with phage infection. The soldiers, represented by the exogenous genes delivered by the “Trojan horse”, effectively function within the cells of *R. solanacearum*.

### Engineered phage delivering CRISPR-AsCas12f1 targets *hrpB* of *R. solanacearum*

The hypersensitive response and pathogenicity (*hrp*) genes encoded type III secretion system, with its delivered type III effectors (T3Es), is one of the major virulence determinants of *R. solanacearum* ^18^. By disrupting the *hrpB* gene, a key regulator of *hrp* genes, namely, a mutant strain of *R. solanacearum* is avirulent and can function as a biocontrol agent against bacterial wilt caused by this pathogen ^19, 20^. We plan to deliver the CRISPR-Cas system that targets *hrpB* into *R. solanacearum* in nature via the engineered phage to disarm the key weapon of *R. solanacearum* and make the pathogen an avirulent biocontrol agent.

To achieve this, a miniature class 2 type V-F CRISPR-Cas from *Acidibacillus sulfuroxidans* (CRISPR-AsCas12f1) ^21, 22^ was selected due to the limited cargo size of the engineered phage. As shown in Figure 6a, the AsCas12f1 gene was placed under the control of a lac promoter, and a 6*his tag was added. The single-guide RNA (sgRNA) was designed with three 20bp spacers targeting 48–67 bp, 642–661bp, and 1210–1229 bp of *hrpB* ORF. Two 400bp homologous arms were designed to edit the *hrpB* targeted region, because non-homologous DNA end joining (NHEJ) is lacking in *R. solanacearum*. AsCas12f1, sgRNA, and homologous arms were cloned to the phasmid vector pRSCqYFP01. However, the cargo was too large to produce an infective phage. The R6kγori sequence (780bp) was removed using the Gibson assembly method for the assembly of three linear DNA fragments amplified by the primers listed in Supplementary Table 3. The Gibson assembly reaction sample was directly transformed into the *R. solanacearum* mutant Δ*hrpB* that lacks target sequences of sgRNA to generate an infective engineered phage, because the resulting DNA without R6kγori cannot replicate in *E. coli*. The resulting infective phage, which now contains the CRISPR-AsCas12f system, was named RSCqCRISPR-Cas (Supplementary Figure 3). An engineered phage without sgRNA and homologous arms, namely, RSCqCas, was also constructed for control.

**Figure 6.**
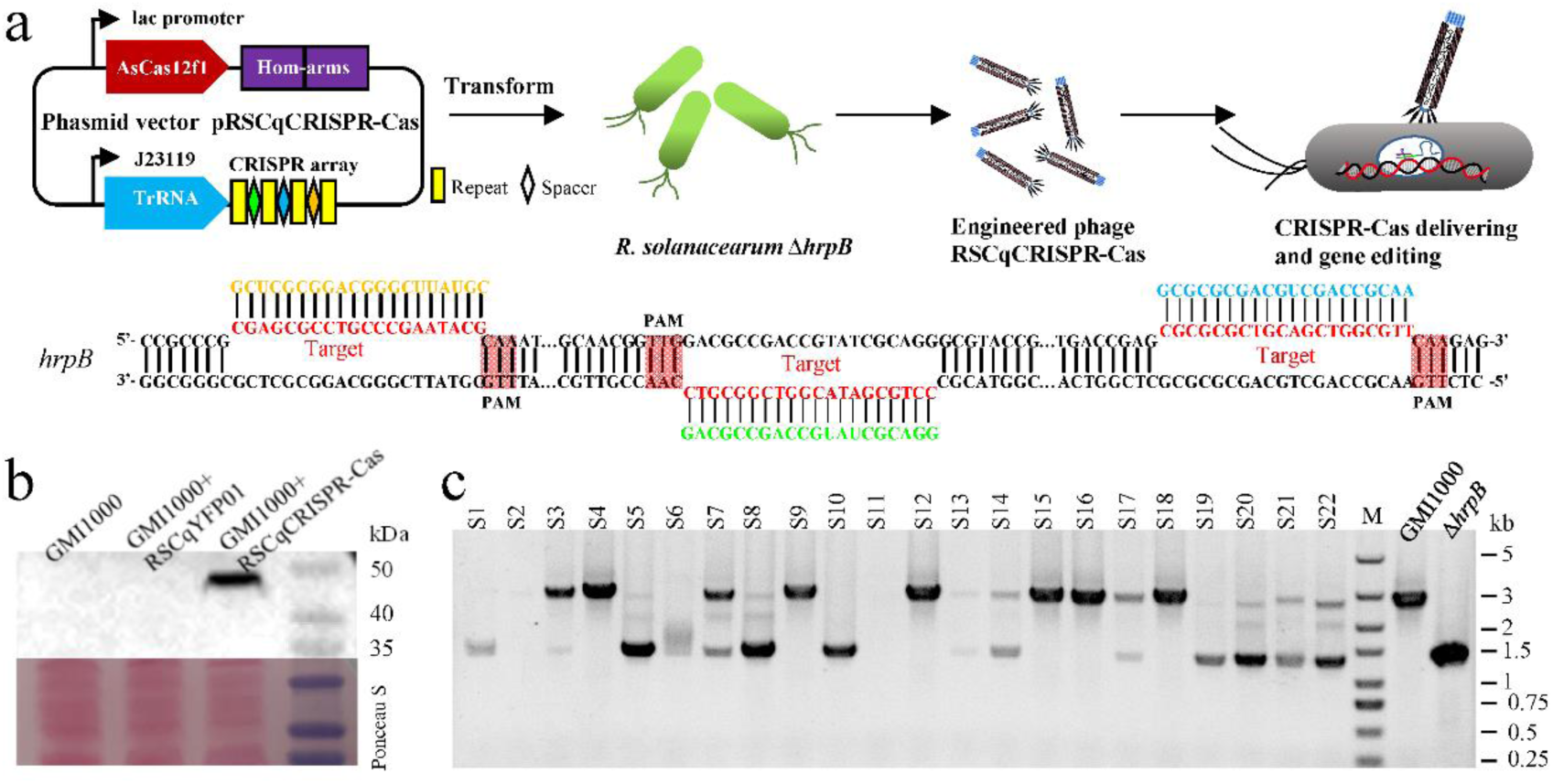
Engineered phage delivering CRISPR-AsCas12f targets *hrpB* of *R. solanacearum*. **a.** Schematic of sgRNA scaffold design and engineered phage RSCqCRISPR-Cas construction. AsCas12f1, sgRNA, and two 400bp homologous arms (Hom-arms) were cloned to the phasmid vector pRSCqYFP01. The R6kγori sequence of the resulting plasmid was removed via Gibson assembly to reduce the size of cargo DNA. The reaction sample of Gibson assembly was transformed into the *R. solanacearum* Δ*hrpB*, generating engineered phage RSCqCRISPR-Cas. Sequences targeted by CRISPR and protospacer adjacent motif (PAM) sequences are indicated. **b**. AsCas12f1 (49.5 kDa expected) in the total protein of *R. solanacearum* GMI1000 infected with engineered phage RSCqCRISPR-Cas was detected via Western blot by using a monoclonal antibody against the 6*His tag. GMI1000 and GMI1000 infected with RSCqYFP01 were set as control groups. **c**. Verification of *hrpB* deletion mediated by engineered phage RSCqCRISPR-Cas via PCR by using primers that bind the flanking sequence of the homologous arms. *R. solanacearum* GMI1000 was infected with RSCqCRISPR-Cas for 48 h in MP medium, plated on BG agar medium, and cultured at 37 ℃. *R. solanacearum* wild-type strain GMI1000 and mutant Δ*hrpB* were used as controls.

*R. solanacearum* GMI1000 was infected with the engineered phage RSCqCRISPR-Cas for 24 h, and Western blot analysis using a monoclonal antibody against the 6*His tag was subsequently performed. As shown in Figure 6b, AsCas12f1, which is expected to be 49.5 kDa, can be detected in the total protein of RSCqCRISPR-Cas infected strain but not in GMI1000 or RSCqYFP01 infected stain, indicating the successful delivery and expression of the AsCas12f1 gene in *R. solanacearum cells*.

To assess the gene-editing effect of the CRISPR-AsCas12f1 system, *R. solanacearum* GMI1000 was infected with the engineered phage RSCqCRISPR-Cas for 48 h in BG and MP media. The resulting culture was diluted and plated on BG agar medium and incubated at 28 ℃ or 37 ℃. The gene editing effect of CRISPR-AsCas12f1 was assessed via PCR by using primers that bind the flanking sequences of the homologous arms. However, no gene editing was detected for RSCqCRISPR-Cas infection in BG medium. When *R. solanacearum* was infected in MP medium and cultured at 28 ℃ after spread plating, most randomly selected colonies showed bands for wild-type and *hrpB* deletion. Wild-type, *hrpB* deleted, hetero-type (showing bands for wild-type and *hrpB* deletion), and potential off-target colonies without any bands were detected when cultured at 37 ℃ (Figure 6c). These findings indicate that the CRISPR-AsCas12f1 system successfully edited the target gene, *hrpB*, although the efficiency and accuracy of gene editing in *R. solanacearum* may not yet be sufficient for gene functional studies.

### Engineered phage RSCqCRISPR-Cas attenuates the virulence of *R. solanacearum*

*R. solanacearum* GMI1000 was infected with the engineered phage RSCqCRISPR-Cas for 12 h in BG medium, and no gene editing was detected by PCR verification after spread plating. The resulting bacterial culture (GMI1000/RSCqCRISPR-Cas) was inoculated to susceptible tomato plants via stem injection to evaluate the effect on the virulence of *R. solanacearum*. As shown in Figures 7a, 7b, and 7c, nearly all the tomato plants inoculated with GMI1000, GMI1000/RSCq, GMI1000/RSCqYFP01, and GMI1000/RSCqCas exhibited wilt symptoms 9 days post-inoculation. However, 96.9% of the tomato plants survived the infection of GMI1000/RSCqCRISPR-Cas, suggesting that RSCqCRISPR-Cas significantly attenuated the virulence of GMI1000.

**Figure 7.**
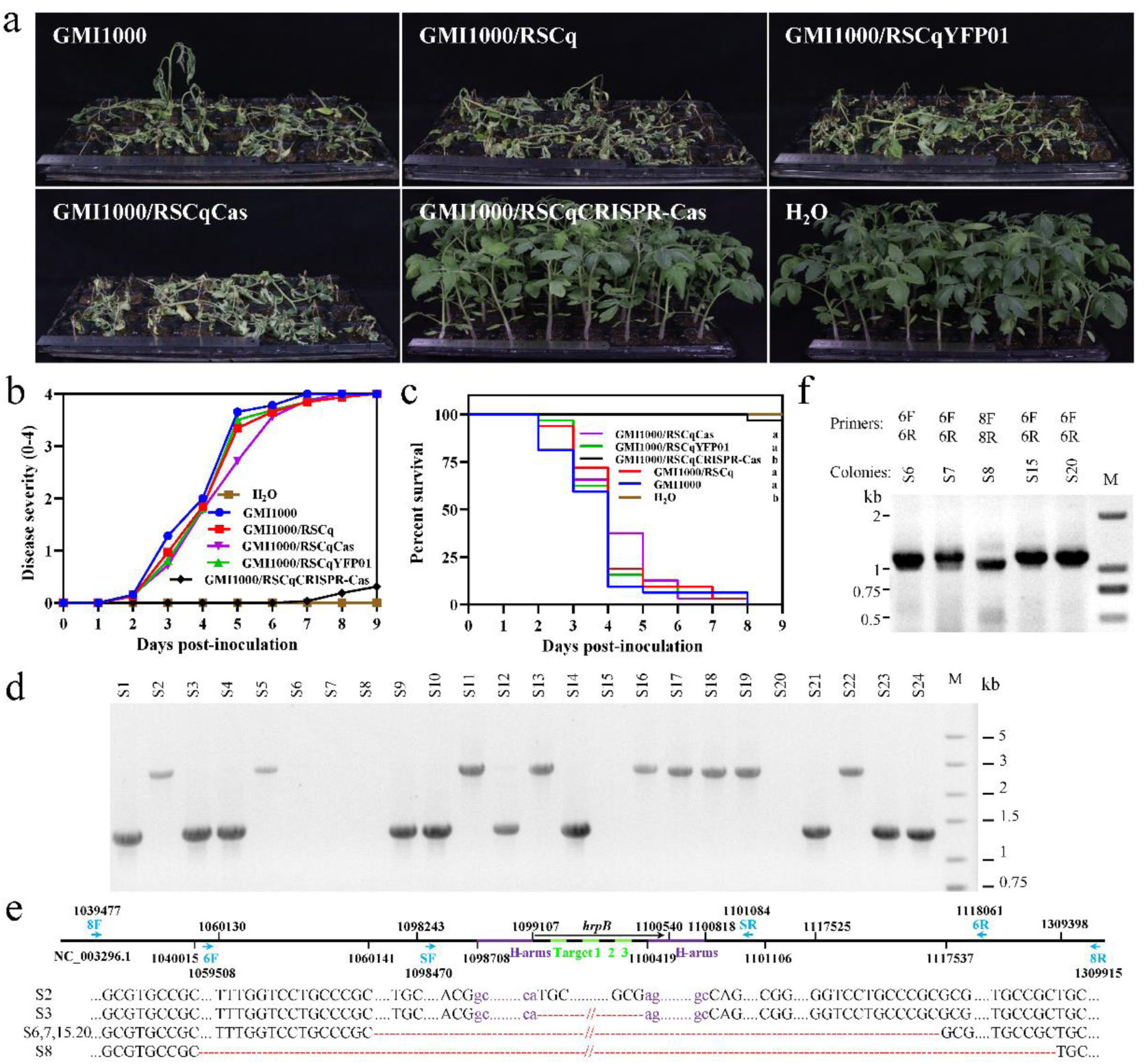
The engineered phage RSCqCRISPR-Cas attenuates the virulence of *R. solanacearum*. **a**. Bacterial wilt symptoms of tomato plants 10 days after inoculation with *R. solanacearum* GMI1000, GMI1000 infected with phage RSCq or engineered phages. Virulence was assayed three times independently, and one representative result was shown. **b**. The disease severity of infected tomato plants was scored on a visual scale of 0 (no symptoms) to 4 (complete wilting) daily. **c**. Survival curve of infected tomato plants. Kaplan-Meier survival analysis with the Gehan-Breslow-Wilcoxon method was used to compare pathogenicity between the mutant and wild-type strains. A P value of 0.05 was considered significant. **d**. PCR verification of *hrpB* deletion mediated by the engineered phage RSCqCRISPR-Cas of colonies recovered from infected tomato plants. **e**. Genome resequencing deduced gene deletion in plant-recovered colonies S2, S3, S6, S7, S8, S15, and S20. The red dashes correspond to deleted nucleotides in the corresponding mutants, while the dots represent conserved nucleotides. **f**. PCR verification of gene deletion in plant-recovered colonies S6, S7, S8, S15, and S20. Primers used are marked in Figure 7e.

*R. solanacearum* was then recovered from the stem of GMI1000/ RSCqCRISPR-Cas infected tomato plants via spread plating. PCR verification was performed on randomly selected colonies to further assay the gene editing effect of CRISPR-Cas12f. As shown in Figure 7d, hetero-type colonies were not detected, unlike in the medium. Among the 24 randomly selected colonies, 10 showed bands indicating *hrpB* deletion, 9 were wild-type strains, and no PCR product was obtained for the remaining 5 colonies. Whole genome sequencing was performed on the 5 colonies that did not yield PCR products, along with a wild-type colony and an *hrpB*-deleted colony. The *hrpB* locus with its flanking sequence was aligned to the assembled genome of the sequenced colonies by using BLASTn. As shown in Figure 7e, genome resequencing confirmed that colonies S2 and S3 were wild-type and *hrpB* mutant, respectively. Moreover, genome re-sequencing found that 57.4 kb of DNA flanking *hrpB* was deleted in colonies S6, S7, S15, and S20. Meanwhile, 269.4 kb of DNA flanking *hrpB* was deleted in colony S8. These deletions were further validated via PCR by using primers that bind to the flanking sequences of the deduced deletion regions (Figure 7f). All these findings indicate that CRISPR-AsCas12f-mediated gene editing persisted during plant infection of *R. solanacearum* and ultimately weakened its virulence on the host plant.

Tobacco (*Nicotiana tabacum*) Yunyan87, a major cultivar planted in China, was selected as the test host plant to confirm the effect of RSCqCRISPR-Cas on *R. solanacearum* virulence. The virulence of RSCqCRISPR-Cas-infected *R. solanacearum* phylotype I Tb04, which was isolated from tobacco in Baise, Guangxi, China ^23^, was determined via stem injection. Similarly, the virulence of *R. solanacearum* Tb04 on tobacco Yunyan87 was significantly attenuated by the infection of RSCqCRISPR-Cas (Supplementary Figure 4).

### Plant bacterial wilt can be efficiently controlled by the engineered phage RSCqCRISPR-Cas

Tobacco Yunyan87 was used as the test host plant to explore the potential application of engineered phage in plant bacterial wilt. The target region of CRISPR-AsCas12f was conserved in the genome of *R. solanacearum* Tb04 (BioProject ID PRJNA616449), which was used as the tested pathogen. Tb04 was watered to soil substrate at a final concentration of 10^8^ CFU/g substrate, followed by engineered phage infection 1 day after *R. solanacearum* watering. The soil substrate was kept wet for 8 days to fully infect *R. solanacearum*. Then 3-week-old tobacco seedlings were transplanted to Tb04 contaminated soil with or without phage treatment. As shown in Figure 8, although phage RSCq and engineered phage RSCqYFP01 treatment delayed bacterial wilt symptoms, the final disease index of tobacco planted in RSCq or RSCqYFP01 treated soil was similar to that of tobacco planted in soil without treatment. However, the survival percentage of tobacco planted in soil with engineered phage RSCqCRISPR-Cas treatment was significantly higher than that of tobacco planted in other soil. RSCqCRISPR-Cas treatment efficiently controlled the bacterial wilt of tobacco with a biocontrol efficiency of 59.2%. That is, the “Trojan horse” viruses that deliver CRISPR-AsCas12f help protect plants from the pathogen *R. solanacearum*.

**Figure 8.**
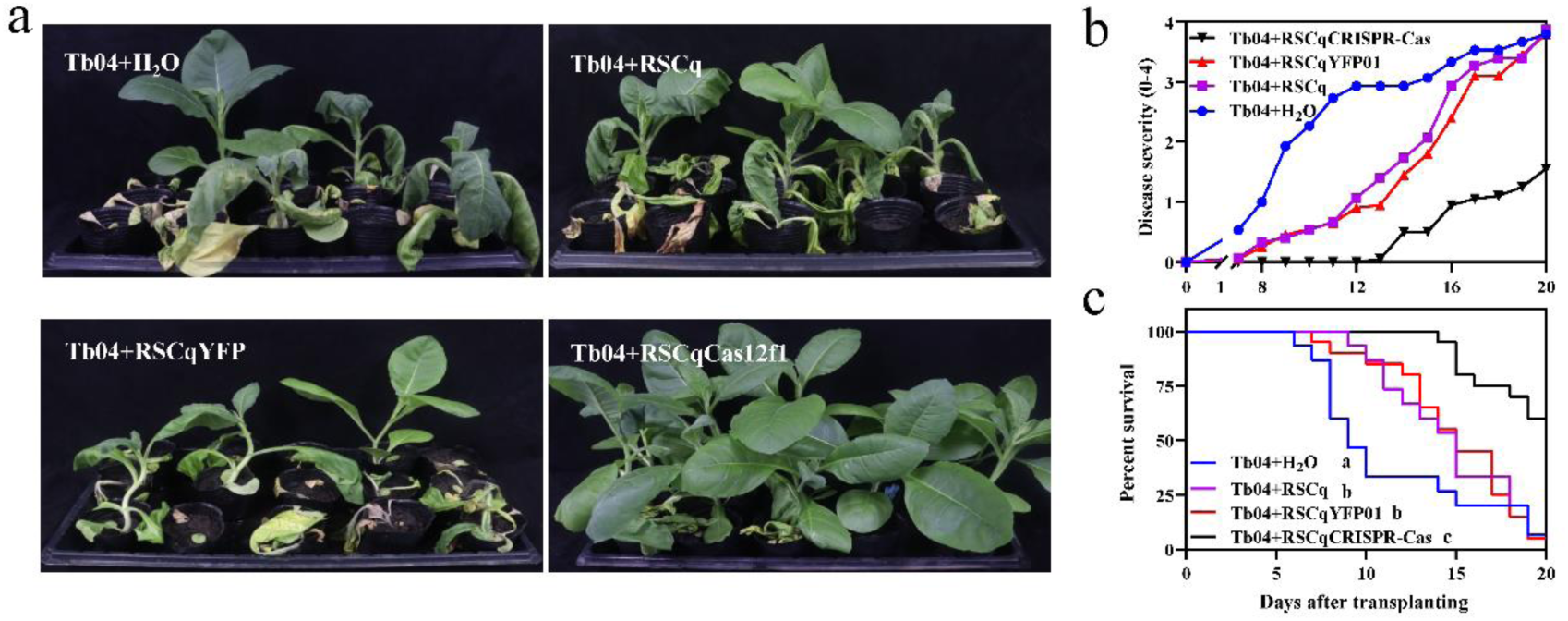
Plant bacterial wilt biocontrol effect of the engineered phage RSCqCRISPR-Cas **a**. Bacterial wilt symptoms of tobacco 20 days after transplants to *R. solanacearum* Tb04 contaminated soils with or without phage treatment. The biocontrol experiment was assayed three times independently, and one representative result was shown. **b**. Disease severity of tobacco was scored on a visual scale of 0 (no symptoms) to 4 (complete wilting) daily. **c**. Survival curve of tobacco planted in soils with or without phage treatment. Kaplan-Meier survival analysis with the Gehan-Breslow-Wilcoxon method was used to compare bacterial wilt of tobacco planted in different soils.

## Methods

### Bacterial strains and growth conditions

Phylotype I *R. solanacearum* GM I1000 ^24^ was mainly used as the model strain. *R. solanacearum* Cq05 (BioProject ID PRJNA974909) and other *R. solanacearum* strains were isolated from Guanxi, China, and are detailed in Supplementary Table 2. *R. solanacearum* strains were cultured at 28 ℃ in BG medium (10 g/L bactopeptone, 1 g/L casamino acids, 1 g/L yeast extract, and 5 g/L glucose) or on BG agar medium ^25^, unless otherwise specified. *Escherichia coli* DH5αλpir was used as the host strain for the replication of phagemid vectors and cultured at 37 °C in LB medium or on LB agar medium. Kanamycin was supplemented when needed at a final concentration of 25 μg/mL.

Prophage sequences in *R. solanacearum* strains with complete genomes were downloaded from the Pre-Calculated Genome in PHASTER^15^. The genome of *R. solanacearum* Cq05 (BioProject ID PRJNA974909) was submitted to PHASTER to predict prophage sequences. Filamentous prophages were determined by “Most Common Phage”, which is an important term in PHASTER defined by the phage(s) with the highest number of proteins most similar to those in the region. The genomic phylogenetic tree of *R. solanacearum* strains with complete genome was constructed via GToTree based on single-copy gene set ^26^ and visualized via tvBOT ^27^.

### Phage isolation and characteristic assay

The filamentous phage RSCq was isolated using a double agar overlay plaque assay. Briefly, bacterial supernatant from the 12 h cultured *R. solanacearum* Cq05 was collected by centrifugation and filtered through a 0.22-μm membrane. The filtered supernatant was then serially diluted to achieve an appropriate plaque count on plates. The bacterial culture of host strain GMI1000 and the diluted supernatant which contains phage RSCq were mixed in 3 ml soft agar BG medium at 50 ℃. The mixture was quickly poured onto a hard agar BG medium surface and cultured at 28 ℃. The single plaque was picked into sterile SM buffer (100 mmol/L NaCl; 8 mmol/L MgSO_4_; 50 mmol/L Tris.HCl; 0.01% Gelatin) for a new round of double agar overlay plaque assay.

The number of plaque forming units (PFU) of filamentous phage RSCq in the supernatant of RSCq infected GMI1000 was determined by double agar overlay plaque assay. The supernatant containing RSCq was added with a multiplicity of infection (MOI) of 10 and co-inoculated with the tested *R. solanacearum* strains. The growth curves of *R. solanacearum* strains with or without RSCq were measured every two hours monitoring bacterial growth (A600) via Bioscreen C Pro (Oy Growth Curves Ab Ltd., Turku, Finland) with three technical replicates and three biological repeats.

The replicative form (RF) DNA of the phage RSCq in the RSCq infected GMI1000 was extracted by alkaline lysis method, followed by phenol extraction. The flanking DNA of the RSCq junction site was amplified by using primers RSCqvF and RSCqvR (Supplementary Table 3) and sequenced by Sanger sequencing. The genome sequence of phage RSCq was determined by the flanking sequence of the junction site and the predicted prophage sequence, and confirmed by whole genome Sanger sequencing.

### Construction of engineered phages

The *eYFP* gene controlled by the lac promoter was inserted between the transposase recognition sequence (ME) and the kanamycin resistance gene (KanR) of the transposon of the EZ-Tn5™ <R6Kγori/KAN-2> Insertion Kit (Lucigen, Wisconsin, USA) via overlap extension PCR by using primers list in Supplementary Table3. The modified transposon was then inserted into RSCq RF DNA randomly in vitro by Tn5 transposase according to the manufacturer’s instructions. The resulting DNAs were transformed to *E. coli* DH5α λpir, generating a transposon-inserted plasmids library. The transposon-inserted plasmid library was electrotransformed into *R. solanacearum* GMI1000, and the engineered phages were screened based on the growth inhibition effect of RSCq on *R. solanacearum* GMI1000.

The *luxA* and *luxB* genes were designed to be under the control of the promoter of the kanamycin resistance gene from the plasmid pK18mobsacB ^28^. The *luxA* and *luxB* gene cassettes were cloned to phagemid vector pRSCqYFP01 at the NdeI and XbaI sites by using the primers listed in Supplementary Table 3. The recombined vectors were electrotransformed into *R. solanacearum* GMI1000, and the engineered filamentous phages RSCqluxA and RSCqluxB were isolated from the supernatant of transformants.

AsCas12f1 and sgRNA scaffold were synthesized at GeneCreate Biological Engineering Co., Ltd, Wuhan, China. Homologous arms were amplified from *R. solanacearum* GMI1000. sgRNA scaffold and homologous arms were fused by overlapping PCR by using primers listed in Supplementary Table 3. The fused DNA fragment and AsCas12f1 gene were cloned to XbaI/NdeI-digested pRSCqYFP01 via Gibson assembly. The resulting plasmid was PCR amplified using primers listed in Supplementary Table 3, generating three DNA fragments to remove R6Kγori. These DNA fragments were assembled via Gibson assembly, followed by transformation to *R. solanacearum* Δ*hrpB*. The engineered phage RSCqCRISPR-Cas was isolated from the supernatant of transformants. The map and the sequence of the engineered phage RSCqCRISPR-Cas RF DNA are available in Supplementary Figure 2 and Supplementary Note 2, respectively.

### Infectivity assay of engineered phage in soil

*R. solanacearum* GMI1000 cultured in BG medium was centrifuged and resuspended in sterile H_2_O at a final concentration of 10^9^ CFU/ml. The resulting culture was mixed with sterile soil substrate at a final concentration of 10^8^ CFU/g substrate. The soil that contains pathogens was treated with the engineered phage RSCqYFP01 at the same volume as GMI1000 one day after GMI1000 treatment. The resulting soil was sampled and spread plated on BG agar medium. The kanamycin resistance of colonies was assayed by replica plating. The phage infection rate was represented by the rate of kanamycin-resistant colonies. This experiment was performed three times independently.

### Luminescence assay of *R. solanacearum* infected by engineered phages

The engineered phages RSCqluxA and RSCqluxB were obtained by centrifugation and filtered through a 0.22-μm membrane from the culture supernatant of infected *R. solanacearum* GMI1000. Phage-free *R. solanacearum* GMI1000 was co-infected by the engineered phages RSCqluxA and RSCqluxB in BG liquid medium for 24 h. The infected bacteria were then subjected to gradient dilution and quantification by colony counting. 1% aldehyde in ethyl alcohol was added to the diluted bacterial culture by 2%, and bioluminescence was detected by using a Synergy II multiplate detection reader immediately. This process was performed with three biological repeats, and the average value ±standard deviation was presented.

GMI1000 infected by RSCqluxA-, RSCqluxB-, or co-infected by RSCqluxA/RSCqluxB were spread plated on BG agar medium after gradient dilution and cultured at 28 ℃. Subsequently, 1% aldehyde in ethyl alcohol was evenly sprayed on the surface of the culture. The resulting plate was immediately imaged under a luminescence imaging system ^29^. Bioluminescence images were inverted (i.e., photographic negatives were generated) and merged with the images taken under white light. This experiment was repeated three times, and representative images were presented. *R. solanacearum* GMI1000 was infected by the engineered phages RSCqluxA and RSCqluxB, generating RSCqluxA/GMI1000 and RSCqluxB/GMI1000, respectively. RSCqluxA/GMI1000 and RSCqluxB/GMI1000 were streaked on BG agar medium crossing each other and cultured at 28℃. The resulting cultures were imaged under a luminescence imaging system as described.

### Western blot analysis

*R. solanacearum* GMI1000 was infected by the engineered phage RSCqCRISPR-Cas for 24 h. The total protein of the resulting culture was extracted by boiling, followed by sodium dodecyl sulfate (SDS) - polyacrylamide gel electrophoresis (PAGE). The target protein was detected by Western blot after transferring the protein from the gel to the membrane. 6*His-tag monoclonal antibody was commercially acquired from Proteintech, Wuhan, China, and was used as primary antibody for Western blot at dilution of 1:10000. HRP-conjugated Rabbit anti-mouse IgG was commercially acquired from Sangon, Shanghai, China and was used as secondary antibody at dilution of 1:10000. The resulting membrane was treated with ECL luminescence reagent (Sangon, Shanghai, China) and imaged under a luminescence imaging system.

### RSCqCRISPR-Cas mediated gene editing assay

Engineered phage RSCqCRISPR-Cas was obtained by centrifugation and filtered through a 0.22-μm membrane from the culture supernatant of infected *R. solanacearum* Δ*hrpB*. *R. solanacearum* wild-type strain GMI1000 was infected by the engineered phages RSCqCRISPR-Cas in BG or MP (MP medium for 1L: 1.25×10^-4^ g FeSO_4_·7H_2_O, 0.5 g (NH_4_)_2_SO_4_, 0.05 g MgSO_4_·7H2O, 3.4 g KH2PO4, 2% glycerol, pH adjusted to 7 with KOH) ^30^ liquid medium for 48 h. The resulting bacterial culture was diluted and plated on BG agar medium, and cultured at 28 ℃ or 37 ℃. Colonies were randomly selected for gene deletion assay via PCR by using primers SF+SR listed in Supplementary Table 3.

Colony samples of *R. solanacearum* recovered from infected tomato plants were subjected to whole genome resequencing via an Illumina NovaSeq 6000 platform at Annoroad Gene Technology, Beijing, China. The sequencing clean reads (deposited at GenBank, BioProject ID PRJNA1012353) were assembled via SPAdes ^31^. The *hrpB* locus and its flanking sequence were used as query and aligned to the resulting scaffolds from genome assembly via BLASTn. The 57.4 kb DNA deletion in colonies S6, S7, S15, and S20 was verified via PCR by using primers 6F+6R listed in Supplementary Table 3. The 269.3 kb DNA deletion in colony S8 was verified via PCR by using primers 8F+8R, which are listed in Supplementary Table 3.

### Pathogenicity phenotyping and biocontrol assay

The pathogenicity assays were conducted following the previously described method ^32^. In brief, the susceptible tomato cultivar Zhongshu No. 4 or tobacco Yunyan87 was cultured in a greenhouse for 4 weeks and used as the test host plant. *R. solanacearum* was cultured in BG medium with the infecting of the engineered phages or the parent phage RSCq for 12 h. *R. solanacearum* without phage infection was used as a control. The resulting culture was adjusted to 10^7^ CFU/mL and inoculated into the stems of 32 tomato plants or 15 tobacco plants by injection. The wilting symptoms of the inoculated plants were recorded daily. Kaplan–Meier survival analysis was performed with the Gehan–Breslow–Wilcoxon method to assay the effect of the engineered phage infection on virulence. Three times of pathogenicity assays were performed, and one representative result was presented.

For the biocontrol assay, *R. solanacearum* strain Tb04 cultured in BG medium was centrifuged and resuspended in sterile H_2_O at a final concentration of 10^9^ CFU/ml. The resulting culture was mixed with soil substrate at a final concentration of 10^8^ CFU/g substrate. The engineered phages RSCqCRISPR-Cas, RSCqYFP01, and the parent phage RSCq were obtained by centrifugation and filtered through a 0.22-μm membrane from the culture supernatant of infected *R. solanacearum* GMI1000. The soil that contains pathogens was treated with the phages RSCqCRISPR-Cas, RSCqYFP01, or RSCq infection at the same volume with Tb04 1 day after Tb04 watering. The soil without phage treatment was set as a control group. Three-week-old tobacco Yunyan87 seedlings were transplanted into the treated soil 8 days post phage treatment. 15 tobacco plants were transplanted for each experimental group. The wilting symptoms of tobacco plants were scored. The control efficiency of phage treatment on bacterial wilt (%) was calculated as (disease index of plants in nontreated soil − disease index of plants in phages-treated soil)/ disease index of plants in nontreated soil × 100. Kaplan–Meier survival analysis was performed with the Gehan–Breslow–Wilcoxon method to assay the effect of engineered phage treatment on plant bacterial wilt. Three times of biocontrol assays were performed, and one representative result was presented.

## Discussion

### More filamentous phages can be efficiently isolated via genome mining

Filamentous phages, regarded as masters of a microbial sharing economy, play crucial roles in promoting bacterial virulence, shaping bacterial communities, and promoting biotechnology developments ^11^. More filamentous phages should be discovered to make the study of filamentous phages flourish. A study used a machine learning approach to mine microbial genomes and metagenomes for inoviruses. A total of 10,295 inovirus-like sequences were found, from which 5,964 distinct species appear to have been identified ^33^. This finding alone represents a 100-fold expansion of the previously described diversity (57 genomes) within the *Inoviridae* family ^33, 34^. These results indicate a vast pool of unexplored filamentous phages that await functional analysis. Our study identified intact filamentous prophage sequences in 42 of the 62 investigated *R. solanacearum* phylotypes I, II, and III genomes, suggesting that filamentous prophages distributed widely throughout *R. solanacearum* phylotypes I, II, and III strains. A filamentous phage was subsequently successfully isolated via genome mining. We believe that more filamentous phages that infect *R. solanacearum* or other bacteria can be efficiently isolated via genome mining, because of the low cost of bacterial genome sequencing.

### Proposed filamentous phage engineering method should be useful

Phage genetic engineering enables deliberate modifications of natural phage isolates to enhance their suitability for various applications ^35^. For example, M13-based engineered phage delivers the CRISPR-Cas system that targets the carbapenem-resistant gene of enterohemorrhagic *E. coli* and significantly improves survival in a *Galleria mellonella* infection model^36^. Staphylococcal phage ΦNM1-based delivers the CRISPR-Cas system that targets antibiotic resistant genes and functions in vivo to kill *S. aureus* in a mouse skin colonization model ^37^. The application of phage-delivered CRISPR-Cas system in agriculture has not yet been reported, and engineering methods for non-model phage are important for this application. Moreover, the aforementioned M13-based and ΦNM1-based engineered phages mentioned cannot be secreted from infected bacteria, and engineered phage particles should be prepackaged with the m13cp helper plasmid or in ΦNM1 capsids. However, engineered phage replication from infected bacteria in nature is important for application in agriculture.

Many sophisticated phage genetic engineering methods have been developed ^38^. One such method is phage recombineering with electroporated DNA. Other approaches to phage modification include the assembly of an engineered phage genome from DNA fragments in vitro, followed by recovery of the engineered phage via the transformation of a suitable bacterial host ^38^. However, the insert site of an exogenous target gene is crucial for in vitro genome assembly. Previous methods relied on comprehensive studies of the functional genomics of specific phages, making them less applicable to non-model phages. Our filamentous phage engineering method proposed in this study followed the aforementioned strategy. We used Tn5 transposase to insert the modified transposon randomly into RSCq RF DNA in vitro, followed by engineered phage recovery. This approach is straightforward to implement and can be employed to genetically engineer other filamentous phages without extensive functional genomic studies.

### Superinfection of *R. solanacearum* is allowed for RSCq

Temperate phages typically encode superinfection exclusion mechanisms to prevent host lysis by virions of the same or similar species. For example, a filamentous phage protein inhibits type IV pili to prevent superinfection of *Pseudomonas aeruginosa* ^39^. In the current study, we show that *R. solanacearum* strain Cq05, which was infected by phage RSCq, can subsequently be infected by the engineered phage RSCqYFP01 and acquire kanamycin resistance. In addition, we observed that the engineered phages RSCqluxA and RSCqluxB can infect a single *R. solanacearum* cell and induce luminescence in the presence of a luminescent substrate. These results strongly suggest that superinfection of *R. solanacearum* is permissible for the phage RSCq. This superinfection mechanism is worth an in-depth study. Moreover, the superinfection makes it possible to deliver larger exogenous DNA overcoming the limitation of cargo into *R. solanacearum* separately by multiple engineered phages.

### More biocontrol factors may be delivered by engineered filamentous phages

The CRISPR-AsCas12f1 gene editing system was used as a therapeutic payload of engineered phages in this study. CRISPR-AsCas12f1 is a novel mini gene editing system that has been recently reported to overcome the limited delivering size of viral-based vectors ^22, 40^. However, *R. solanacearum* gene editing mediated by engineered phage-delivered CRISPR-AsCas12f1 was modestly detected when culturing in minimal medium MP, but was not detected when culturing in rich medium BG. The gene editing efficiency and accuracy of CRISPR-AsCas12f1 in *R. solanacearum* should be optimized further. Recently, an engineered hypercompact CRISPR-Cas12f system based on AsCas12f has been demonstrated to be 11.3-fold more potent than parent protein ^41^. Engineered AsCas12f and other miniature gene editing systems, such as Cas12n ^42^, can be potential plant bacterial disease biocontrol factors delivered by engineered filamentous phage. Further testing is necessary to evaluate their biocontrol effects.

Using resistant cultivars is considered the most effective strategy for controlling bacterial wilt and other plant diseases ^43, 44^. However, the *R. solanacearum* species complex is highly heterogeneous in nature, posing challenges to developing crop disease resistance. For example, in tomatoes, the polygenic resistance to bacterial wilt in the resistant cultivar Hawaii7996 was suggested to be strain-specific ^45^. Type III secreted effectors of *R. solanacearum* play key roles in crop disease resistance, and the diversity of type III secreted effectors in *R. solanacearum* species complex significantly impeded disease resistance breeding effort ^46, 47^. The gene delivery strategy employed in this study can be applied to regulate the effectome of *R. solanacearum* species complex in nature to facilitate disease resistance breeding. For example, certain avirulence genes recognized by the resistance gene of crop cultivars can be delivered to *R. solanacearum* by an engineered filamentous phage, thereby converting the virulent *R. solanacearum* strain to an avirulent strain. However, many effectors are avirulent and virulent dual-functional. The potential risks of delivered avirulence genes should be fully evaluated.

### Biocontrol effect in a field with a long timescale should be further studied

We demonstrated that engineered phage that delivers CRISPR-AsCas12f controlled bacterial wilt of tomato and tobacco plants efficiently in a greenhouse. However, the field environment is more complex than a controlled greenhouse setting. Therefore, the biocontrol effect of the engineered phage on bacterial wilt in field conditions remains uncertain and requires further investigation. Bacterial resistance to phage infection is a significant obstacle in the widespread application of lytic phage reagents. By contrast, filamentous phages establish cooperative relationships with their bacterial hosts and exert minimal burden on them. However, interaction studies between filamentous phages and their host bacteria remain limited, and the occurrence pattern of bacterial resistance to filamentous phages is still unclear. Studying the long-term biocontrol effect of engineered phages is also an important area for future research.

Notably, the World Economic Forum has awarded designer phages as one of the Top 10 Emerging Technologies of 2023. Engineered phage therapeutics in animal bacterial infections have been widely studied and have shown their feasibility^35, 36, 48, 49^. However, engineered phage biocontrol in plant diseases has been largely overlooked. Our study demonstrated the efficient control of tomato and tobacco bacterial wilt by using engineered phages. The proposed phage engineering method is universal and suitable for non-model phages, and we believe the biocontrol strategy in this study can also be applied to other plant bacterial diseases. Further research in this area holds considerable promise.

## Acknowledgments

This work was supported by the National Natural Science Foundation of China (Project 32260713) and the Natural Science Foundation of Guangxi (2023GXNSFBA026208).

## Author contributions

D.Z. designed the research. S.P., H.Q., F.N., F.S., and D.Z. performed experiments. D.Z. wrote the manuscript with input from the other authors. S.P., Y.X., and D.Z. collected and analyzed data. Y.X, L.R., and D.Z. revised the manuscript.

## Competing interests

The authors declare no competing interests.

## Notes

### Competing Interest Statement

The authors have declared no competing interest.

