## Supplementary Figure for "Trojan horse virus delivering CRISPR-AsCas12f1 controls plant bacterial wilt caused by *Ralstonia solanacearum*"

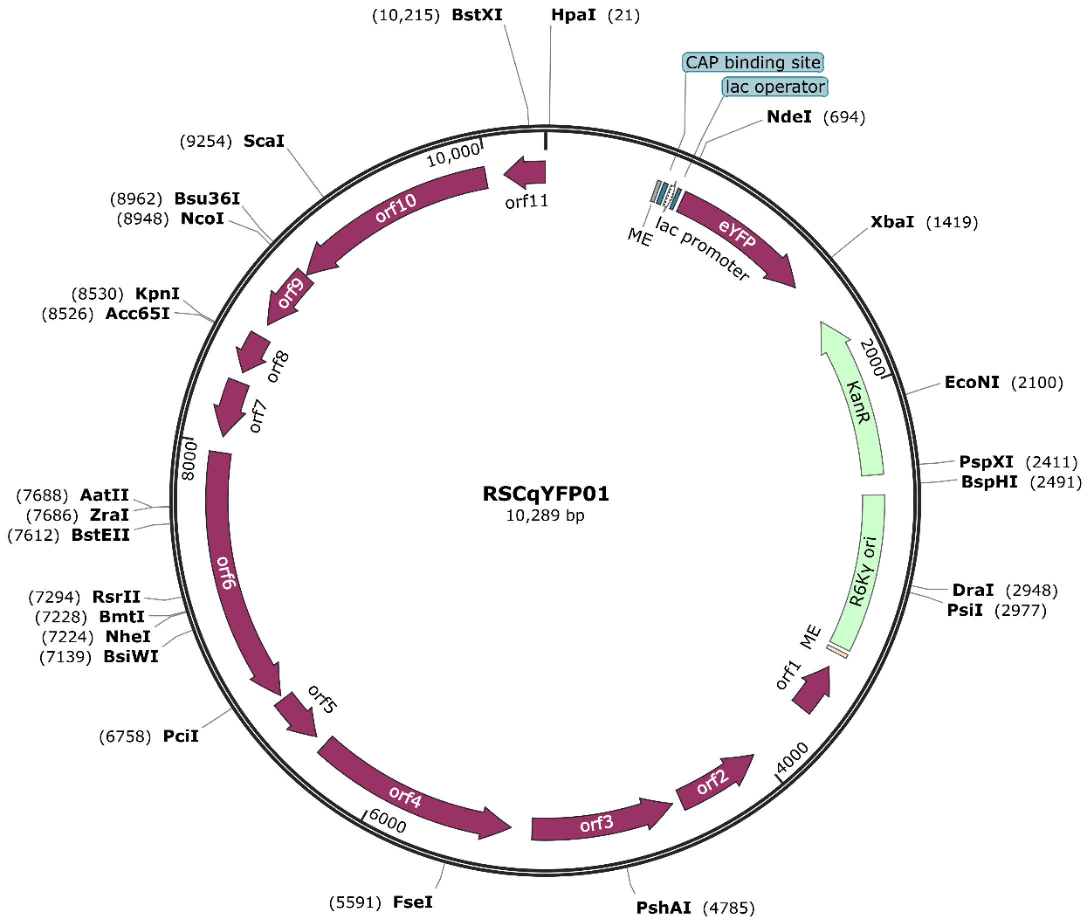


**Supplementary Figure 1**. The replicative form DNA map of the engineered phage RSCqYFP01

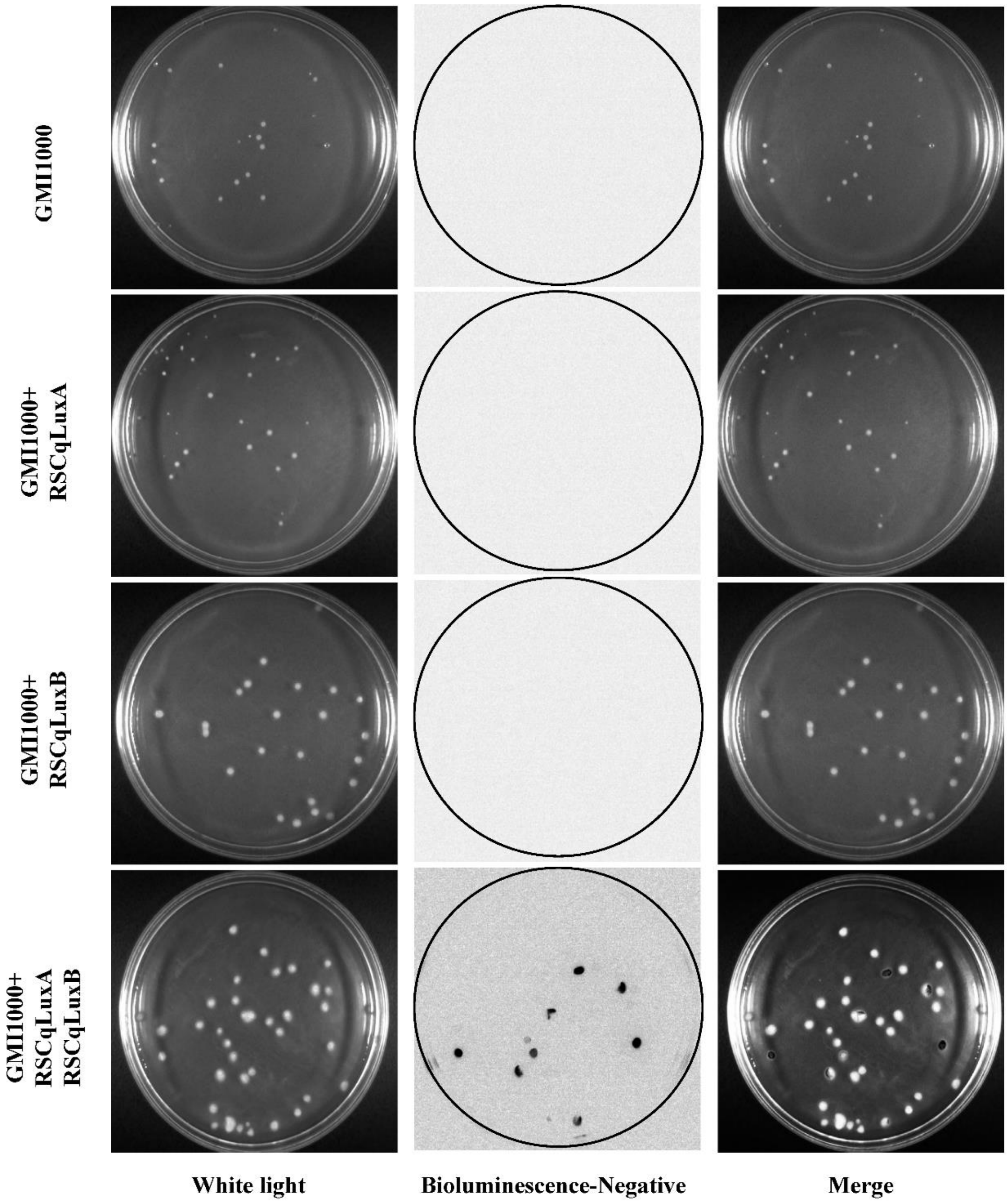


**Supplementary Figure 2**. Luminescence imaging of the engineered filamentous phages RSCqluxA- or RSCqluxB- infected, or RSCqluxA/RSCqluxB co-infected *R. solanacearum* GMI1000 spread plated on BG medium. Left panel, image of RSCqluxA/GMI1000 and RSCqluxB/GMI1000 under white light. Middle panel, the negative image of bioluminescence in the dark. Right panel, merged image.

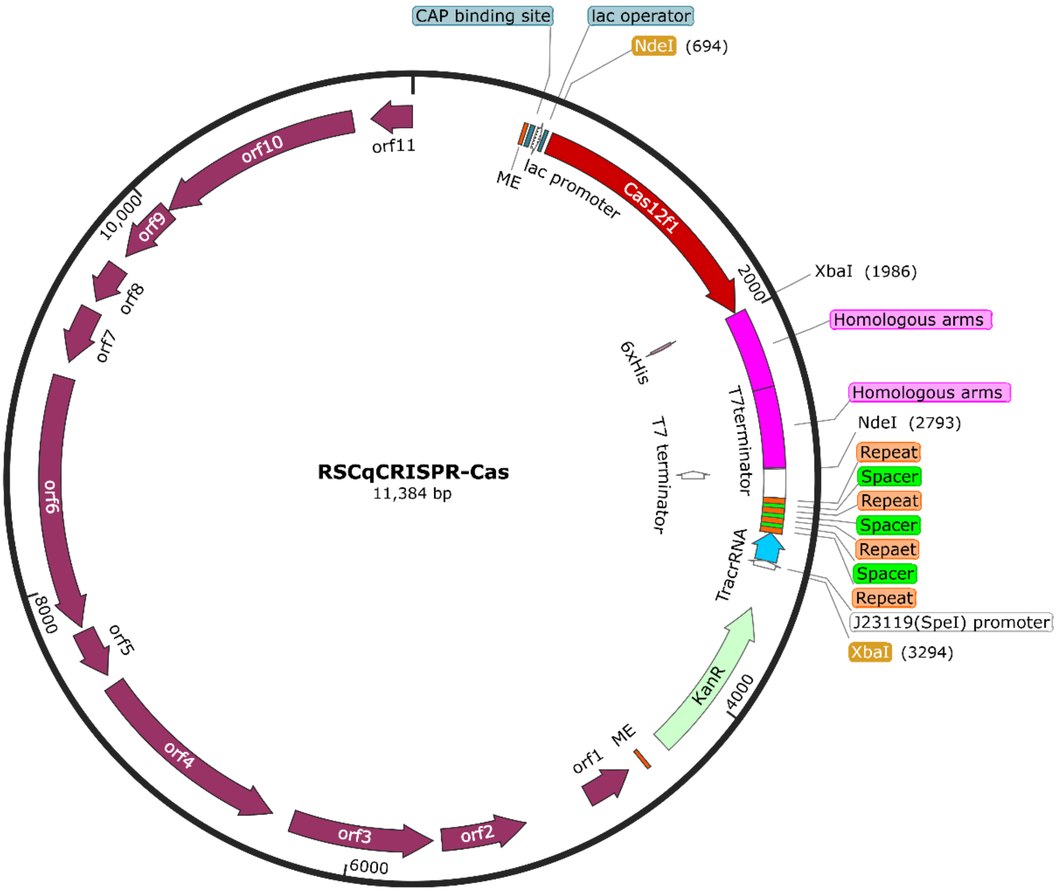


**Supplementary Figure 3**. The replicative form DNA map of the engineered phage RSCqCRISPR-Cas

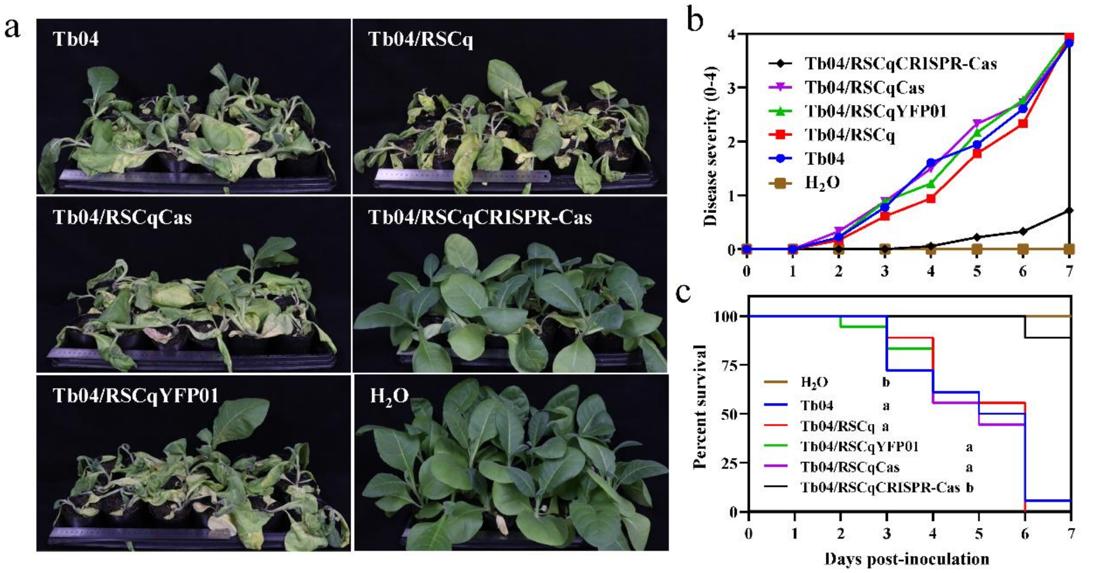


**Supplementary Figure 4**. Virulence assay of engineered phages infected *R. solanacearum* Tb04 on tobacco Yunyan87. **a**. Bacterial wilt symptoms of tobacco plants seven days after inoculation with *R. solanacearum* Tb04, Tb04 infected with phage RSCq or engineered phages. **b**. The disease severity of infected tomato plants was scored on a visual scale of 0 (no symptoms) to 4 (complete wilting) daily. **c**. Survival curve of infected tomato plants. Kaplan-Meier survival analysis with the Gehan-Breslow-Wilcoxon method was used to compare pathogenicity between the mutant and wild-type strains. A P value of 0.05 was considered significant.

**Supplementary Note 1**. The whole genome sequence of the engineered phage RSCqYFP01

cgtttacccctcgcccgggttaaccggctatatttacgccagaatcacgggcgtaattcatcacggacataatattaattacatccttcatgccgtc
aagcaaagaggtgcttagatgagaattgagaaatacttagatcaggcgatcgaacgccacggcctgaagaacgacagcaagctggcagag
atgctaggtgtggtgcaaagcgcggtcagccactaccgcaccggccgccgcacggcggacaacgaagtgtgcctccgcctggcgcagct
gctcgagatggagaacccgctgccgatcatcatggcggccgacatggaccgcgccgaacgtgctggccagcactctctctgggaagtttttt
cgacgaggatggcagccagtaacgcgacagccgccctcctcctggtactggtcgcgagcgcaacaaattttgttgcgccctctcccgccaaa
gccgcgccgttgagccattcgacagctcaacgattattgttatgtaaaatagctcgccgacttcgggagcgcctacagcaagcgctgcgtgca
ctgtctcttatacacatctcgcgcaacgcaattaatgtgagttagctcactcattaggcaccccaggctttacactttatgcttccggctcgtatgtt
gtgtggaattgtgagcggataacaatttcacacaggaaacacatatgatggtgagcaagggcgaggagctgttcaccggggtggtgcccatc
ctggtcgagctggacggcgacgtaaacggccacaagttcagcgtgtccggcgagggcgagggcgatgccacctacggcaagctgaccct
gaagttcatctgcaccaccggcaagctgcccgtgccctggcccaccctcgtgaccaccttcggctacggcctgcagtgcttcgcccgctacc
ccgaccacatgaagcagcacgacttcttcaagtccgccatgcccgaaggctacgtccaggagcgcaccatcttcttcaaggacgacggcaa
ctacaagacccgcgccgaggtgaagttcgagggcgacaccctggtgaaccgcatcgagctgaagggcatcgacttcaaggaggacggca
acatcctggggcacaagctggagtacaactacaacagccacaacgtctatatcatggccgacaagcagaagaacggcatcaaggtgaactt
caagatccgccacaacatcgaggacggcagcgtgcagctcgccgaccactaccagcagaacacccccatcggcgacggccccgtgctgc
tgcccgacaaccactacctgagctaccagtccgccctgagcaaagaccccaacgagaagcgcgatcacatggtcctgctggagttcgtgac
cgccgccgggatcactctcggcatggacgagctgtacaagtaatctagaccgccacggttgatgagagctttgttgtaggtggaccagttggt
gattttgaacttttgctttgccacggaacggtctgcgttgtcgggaagatgcgtgatctgatccttcaactcagcaaaagttcgatttattcaacaa
agccgccgtcccgtcaagtcagcgtaatgctctgccagtgttacaaccaattaaccaattctgattagaaaaactcatcgagcatcaaatgaaa
ctgcaatttattcatatcaggattatcaataccatatttttgaaaaagccgtttctgtaatgaaggagaaaactcaccgaggcagttccataggatg
gcaagatcctggtatcggtctgcgattccgactcgtccaacatcaatacaacctattaatttcccctcgtcaaaaataaggttatcaagtgagaaa
tcaccatgagtgacgactgaatccggtgagaatggcaaaagtttatgcatttctttccagacttgttcaacaggccagccattacgctcgtcatca
aaatcactcgcatcaaccaaaccgttattcattcgtgattgcgcctgagcgagacgaaatacgcgatcgctgttaaaaggacaattacaaacag
gaatcgaatgcaaccggcgcaggaacactgccagcgcatcaacaatattttcacctgaatcaggatattcttctaatacctggaatgctgttttc
cggggatcgcagtggtgagtaaccatgcatcatcaggagtacggataaaatgcttgatggtcggaagaggcataaattccgtcagccagttta
gtctgaccatctcatctgtaacatcattggcaacgctacctttgccatgtttcagaaacaactctggcgcatcgggcttcccatacaatcgatagat
tgtcgcacctgattgcccgacattatcgcgagcccatttatacccatataaatcagcatccatgttggaatttaatcgcggcctcgagcaagacgt
ttcccgttgaatatggctcataacaccccttgtattactgtttatgtaagcagacagttttattgttcatgatgatatatttttatcttgtgcaatgtaacat
cagagattttgagacaccctgtgaatgcgcaaaccaacccttggcagaacatatccatcgcgtccgccatctccagcagccgcacgcggcgc
atctcgggcagcgttgggtcctggccacgggtgcgcatgatcgtgctcctgtcgttgaggacccggctaggctggcggggttgccttactggt
tagcagaatgaatcaccgatacgcgagcgaacgtgaagcgactgctgctgcaaaacgtctgcgacctgagcaacaacatgaatggtcttcg
gtttccgtgtttcgtaaagtctggaaacgcggaagtcagcgccctgcaccattatgttccggatctatgtcgggtgcggagaaagaggtaatga
aatggcagatccctggcttgttgtccacaaccgttaaaccttaaaagctttaaaagccttatatattcttttttttcttataaaacttaaaaccttagag
gctatttaagttgctgatttatattaattttattgttcaaacatgagagcttagtacgtgaaacatgagagcttagtacgttagccatgagagcttagt
acgttagccatgagggtttagttcgttaaacatgagagcttagtacgttaaacatgagagcttagtacgtgaaacatgagagcttagtacgtacta
tcaacaggttgaactgctgatcttcggatctatgtcgggtgcggagaaagaggtaatgaaatggcatccggatctgcatcgcaggatgctgct
ggctaccctgtggaacacctacatctgtattaacgaagcaattcatcgatgatggttgagatgtgtataagagacaggtccaaacaagcccgaa
aacggcaccgtgctgatcgacagcacagcagcagatctagcacctcccgttccgaaacgcctacagcaagcgctgcgtgcagtccaaaca
agcccgaaaacggcaccgtgctgatcgacagcacagcagcagatctagcacctcccgttccgaaacgcctgcatctagcaggcgttttttttt
cgtgcggatctttcgatagtaccgatgccgagcggcatcctggtagccccggcgcagtgtccggatatgccagtacagatgcctgatctcgc
gcgccacggccttcagttcgtcgtctgacactgccgattttggttgttctgagcagcgtagcatcggcacgcgaagcgatccgccacagccag

cgcaaccgaagatcgtaagcgtttcgaaagctggcttgcaggaaagggggtgtttgtatgtgggcggccctgtatgtacgtacctggtcatac
ccgctgctcggtcccgccaggcgccctccgggcatcaaccgtcctgcacctggtcaactctcccatgtccccagaggggccccctagcggc
tccgactacggcgcggacactcccgacgctggacgcaaaaaaaggccggcgatccgcacggcctttctatttttcgttaccttttcgttactcc
gggtagaactccggcgacagcacctgctcgggtatccacggccactgctcggggaacacgtccaggccagtctcgtccacagccgccgag
accgcatccgcccagatctcttccttccaatccggatcagccaacatcggctgcaggctgggcgtccgatgcaaccgcgccagaattgcgcg
acgctgttccttgatcgtgcgctgccagctcgaaccccggcggcccggctgatattgccacttgagcaaatgcgccagcagtaccgccatgc
ggcttgccaactcccgctgttcgctcttgcccacgtcctcaatctcctccgcgatgtgccgaatgtcgatgtctgaaagcttgccggcccgcaa
cagcgccgcctgctcactggcccacgccaccacgtccacttcgtagctcgttcccataacgcgctccctccaataaccggatcattttagacgt
ttcgcgttacgccttgcgcgtaacgactttgcgatcaccataaattgcctcgccagccgcttgcatctgcgcctcactcacggccacagaacgg
acaaacgaagcgcgccacaactcggactcccgcaccttctggcgctgttcgaacgtattcgccttcgcagccaacgcgagcaagtccagcat
tgcgtcgaacgaaacgggatttccgttactcggcgtaacggatttttgctgcgactcaaacatcgcaaccttctcttccaagacgccaacataac
gcaacgcttcctgacgcccgtcctcgcacgtcataagctgccggcgcaacgcctcaatctccgcaagatttccgttactcagcgtaacggattt
ggcggcatcgctgatctcttccctcagccgcctattctcagcaaacgccttattgcgttcagcctccgccagctccacctggcgacgcatgctc
accagatcagcgcgaaacttctcgacctccaggcggcattcatcgtaagcatccacgtcggaaagcatttccgttacggtcatacgcttagtaa
cggatttagcctcaacaggctgctgagccttacggcgagctcgatatgcggcttgacgctccgcgttggtcatagcgtgcgccttacgcggac
gaccgcggccacgcctttgctccatccccggcagatcgaccgtcacattatccgttacgtcacgcatattcggctcccgatttgatgattccattt
tacgttactggtaacggcaattcaattatcgttacaatttagttcgtaacggaaatttcgcgagcagccccggcgctagacgccctcagcgggct
ttggagccgtttttctgactgggcggtcgcaggggtcagctgcgccgaaattggctcctgcgccgcttgtggctcgcggtaggggtcaaacg
gcggcctcctgatccactcccgacactgcatgtcatcgaggccggcatccgtgccttgagccgtgtagcacgtgcacctggtcgacgtgcag
gcaccgccgatcaccgtcggcatcgagcgaatctggcgcaactgcgcataggccggcgccgtctcaggccggccagagacagcaggga
cgaacgccgcgaggatctccgccccgtccgtcgccgcggcggacggacccgccttgaccgctaccgctagagctgaagccacaccgccc
ctgccctgctcggccgcaacaggcttagccggagtcgccgtcgtcaactggccggtacggtaatagacccggtacgccaagaacgccgcg
atcaacacgcaggcgatgaacagcatcagcaccggcggtacggtgtacttgcgcttgatgtgcagactggaggacttgtacaggccgaaac
tggacttcggcaggctccacttcttcttgatcggcgcggtgttgaacgtctccgggttcgcgcactccggccattcgtagtaccagcgtccgag
caagccagcgtcccgcaggtggacatgctgccccaccagcttgcggatatgactgtccaagaacgtcgggttctgcgtgatcagaacaaac
gtcacgccggtatgccgcaccgtctcaaacgccgccacgtggtcaggcaccttggacccggccgtgcggacgcgaaacacccgctgcgc
ctcatccaacacgatcagcgagttcggcgggaaggtgaagtacggcagcatcatccctgggttttcagggtcttcgcgcagctccgtccagtc
cgaaaccgccggcgtcgggatatacggcagcttcagctccgggatgcccatgacgaagagcggacgcccctggtcgacggctgccttcat
catctggaccgccaacgcggtcttcccgccaccaggcgtggccgtgatcagcgtgatcggttgcgttgcgctcatgtcagcttgcccagccg
cttcagagtgatcatggagatgcgcgcggtgatgccgccggcgatgatcgacagcccggtgaagacaccgccccgcgccaggatggccg
cagcaacggcaggcatgccggccaggctactcttggccgcgcccagggcagcgctcaccgccgcatccaacccgacataggtgatcaga
ccgatcccgagcgacaccagtagctgacgcgcgagtggcccaacgagggccatgaggaacccagcgagcggcatcactcctcccctttac
ggcctacgccgatgacgatcagcgcagcgcccaaccatgcgcacgcgatgatgaccggtcgaaacatgtcagcaccgtcacagaccggtt
tgagcgaccaagagatcggcatgccgtggatggaggcggtcagatcggacggacacggtgcggtatctgcgccccaaccgctgtccggc
atgaccttgacgttgacctgctgctccttcagatcgggcccgtccggtatctcgccttgctcgatacaacccatgcgcgtctcatggcccgagc
actggtcgtcctgccgctccggagccttgccgttaccagtcgccggatccgtcgccggattgccgttcgcatccacctcctgcttagcggtcgt
caacgtggctgtcttgccatcggaatttggcgtgaccgtagcaacatcgcggtaacgcttacctgtaacgggatcaacgtacgggtcgctcag
attgacgttgactggagtggtggacggcgtgagcttcaccggaatcggcaccttggctgcggccatatcgctagcaacagcagcgggcagc
ggatacgtcaaacctttgttccaatccgcatcgcttgcgcccaccgtcggaccggcagggtcgggcacacatgccgagccactcacgacata
gccatcaacgcagctggacgcctgactcgtgcccgcatagaaattgtcgccgccatcgttggtgtaatggcactcatagctcgtcccattaccg
gtagctttcatacccgcgaacttagccttctgcccagccaaatacgcgtctgccgccaacatcgcagagcacgcggcgccaggagaagcag
caacgccattcgcgatattgccgccggtggcggacgtgttgtagccgtaattccagccccagccattgaaccccgtgtcgccagcagccggt
gaccgcttcgatgtgcaccacgtaccgtccaggcacttctggatgccgagctgcgccaagtaggccaacgacgtcgcggtcgcgatcgcag
gcgtcgcacgaagcgcggccagcgcaacggcagcagcgccttcgcccaccgtcatcgttgcggcgatcgtggcggcacctgcgccaacc

gcggcacgcatctcggtagccgtcgcaacttcagacagcgtcacagcaccagtcgtgacatacccagtccccgtcaacacgatgttcggcg
gcggaatcaacgggatggtcgaagcccacgcggacgacgcccaccagcagagcagcagcacgagcacgcgcatcacagccccctgaa
aatgattactgccggcagaaccggcagcaggaatccggcccagagccagaaatcgatcgcgagcatcacagccccttttcatcaccacca
gcgcccaggccgccaccatcgcggcaacgacgccccaccccatcgtcatgccatccttgaagctctcttgcgggtcacatgccggaaacga
cagcgacggcaacgacgcatcggtcaacgtgccgacgctcccatcactgcccacctgataccgccgcagcacccacccgcccgtggtctt
gacgaactcagacagataggtgactgccccaggtgtctgcgacggagcgacggcgctgtaataggcatccgtggccatgcccacatcggc
aaagcaccgtgcgcccaccaacgcgccgtcagcagccatcacacgctccgacgcatgtacttgaacgtcgcaatgccgatcacgatgatca
gcgccaggcccgccagcgtggcggtgtccgccttcgaatcggacatggcggtagcaacgtcggtcggcaccgcagccatcgccgaaccg
gccagtgcaacggtacccgcggcaacagcagccgccttgctcttgatgctcttgaacatggttttctccttgagatggaggttgagaaaagctc
cgggccgttcaactcccagagccagggaacatcacgccttcttgggctcagcctcagcacgctgcagcggcttgatgctcgtaacgaccttct
gaccgcccttatccttgccgttgctcgtctcgaccatcgagacttccgcgatgaacgggaacgggttgtggatgatcgccttcacaaccgcag
agctctcgcacttcagctcctgcgtgcaggtgcccttcgagtcttcgccacgcagctccacatccgtgtagatcttcccggtgtccagctgcttg
ccatccatgttgccaacccacgtcttggcgccccggatggtcacgcgtgcaatcatttccatggtttcactcctcaggtttcggcactggatcgt
gcgtgccgtacacgtgcctcgccagggcggatttgtgcagctgagccggtacgccctgacggcgtatcgcgacaaccagcgcagcgatgt
cttcctctgtgcagcgcagctcgtaatcgaccgtagggccgaactgcgtctggatgtgcttgagcttgcgttcgcggatggtctcgtcctgaag
ctccaaggccttgacctggtcagtaggaacgcgctgcggatccgcagccatgaaggcctccagggccttgtacgcaccagcgaagtactgg
tcgcgcttgataaggatttcgtgcggaatcacgcgatccttggcgccgaactcgatctcgagacgcacccactcgctatcctgattgccgagct
ggcggcccttctcgtaagcccgcagcatcttgccgttcgcccgacggccaatctcgaacgtcgtaccgcgacagcccttgctgcccgccacg
ccgctctcgatcttgcgatacgtcgggatacgcccgcccgcgttgaagtcgccggcgtagtacagctcttccatctgcgcgatgctcacctcg
ccctggcagaagtccatcgccaggtcgcaccgcgtgatccgcgcgtcgaggtcctgcaccatcgcgtagacggcttgccagtcgccaatcg
cggtgcagccctgccccggccagtccaccaggatcgtgccgccaacgtgctcgccgccgcaggcaacgatcccgagcttcatcgtctcgc
cgttgatgaacgccagcaggtcgtagctgaactcataccggcggaaccctttgcccgcaggcttcatcgtcaccggcaccgagaacaccag
ctggaagtacctgcgcagctgctccagggcgtcgctgatgctgccgtcgggcaggaacgtgaacttgaaccagtccacgatcgcacctgcc
ttgcgttctggactttccccgggttttaccggacccggggaacggcctgtcggccgcccagcctcgctccgctcggcccggtcggccgccat
gccgttctcgcgtgcgagcgccggcccatgaactccatacgcaatgcctgtgggatcaagatgacgagagggggtgggacggacgcgcat
cagcgtgcgccctcgcagcgataccagttcgccagatcggcctggcgcgagttcttcaggaggtcgtcaatgcgatgcatcgcaggacgcg
attgcaggcagcaggcgccgcgcagaacggcagccatcacgatggccagctgcgcaccgtgcgggtgaagcacgtcgccaatcaggtcc
agctgcctatggtctctctgctgcat

**Supplementary Note** 2. The whole genome sequence of the engineered phage RSCqCRISPR-Cas

cgtttacccctcgcccgggttaaccggctatatttacgccagaatcacgggcgtaattcatcacggacataatattaattacat
ccttcatgccgtcaagcaaagaggtgcttagatgagaattgagaaatacttagatcaggcgatcgaacgccacggcctgaa
gaacgacagcaagctggcagagatgctaggtgtggtgcaaagcgcggtcagccactaccgcaccggccgccgcacgg
cggacaacgaagtgtgcctccgcctggcgcagctgctcgagatggagaacccgctgccgatcatcatggcggccgacat
ggaccgcgccgaacgtgctggccagcactctctctgggaagttttttcgacgaggatggcagccagtaacgcgacagcc
gccctcctcctggtactggtcgcgagcgcaacaaattttgttgcgccctctcccgccaaagccgcgccgttgagccattcga
cagctcaacgattattgttatgtaaaatagctcgccgacttcgggagcgcctacagcaagcgctgcgtgcactgtctcttata
cacatctcgcgcaacgcaattaatgtgagttagctcactcattaggcaccccaggctttacactttatgcttccggctcgtatgt
tgtgtggaattgtgagcggataacaatttcacacaggaaacacatatgatgatcaaagtttaccgttatgaaatcgttaaaccg
ctggacctggattggaaagaattcggcaccatcctgcgtcagctgcagcaggaaacccgcttcgcgctgaacaaagcaac
ccagctggcgtgggaatggatgggtttcagctctgattacaaagacaaccatggcgaatacccgaaaagcaaagacatcc
tgggctacaccaacgttcacggttacgcataccacaccatcaaaaccaaagcttatcgtctgaacagcggcaacctgagcc
agaccatcaaacgtgcgaccgatcgcttcaaagcgtaccagaaagaaatcctgcgtggtgacatgtccatcccgtcttaca
aacgtgatatcccgctggatctgatcaaagaaaacatcagcgttaaccgtatgaaccacggtgattatatcgcttccctgtcc
ctgctgtctaacccggcgaaacaggaaatgaacgttaaacgtaaaatctccgtgattatcatcgttcgtggcgcgggtaaaa
ccatcatggatcgcatcctgagcggcgaataccaggtgagcgcgagccagatcatccacgacgaccgtaaaaacaaatg
gtacctgaacatctcttatgatttcgaaccgcagacccgtgttctggatctgaacaaaatcatgggcatcgatctgggtgttgc
ggtggcggtgtacatggcgttccagcacaccccggcgcgctacaaactggaaggcggtgaaatcgaaaacttccgtcgtc
aggtggagtctcgtcgcatctccatgctgcgccagggtaaatacgcgggtggtgctcgtggcggtcacggccgtgataaa
cgtatcaaaccgatcgaacagctgcgtgacaaaatcgctaacttccgcgataccactaaccaccgttactctcgttacatcgt
tgatatggctatcaaagaaggttgcggtaccattcagatggaagatctgaccaacatccgtgatatcggctctcgtttcctgca
gaactggacctactacgatctgcagcagaaaatcatctacaaagcggaagaagcgggcatcaaagttatcaaaattgatcc
gcagtacaccagccagcgttgcagcgaatgcggcaacatcgattccggtaaccgtatcggtcaggcgatcttcaaatgcc
gtgcttgcggctacgaagcgaacgcggactacaacgcagctcgtaacatcgcgatcccgaacatcgataaaatcatcgcg
gaaagcatcaaacatcatcatcatcaccattaatctagagccgtgaagctcgccgcgcccgtgatgatgacgatgctgctga
tcgatgccggcatcggcttgctggcgcgcgcggcggacaaactcgagcccacctcgctcggccagccgatcaagggcg
cggtggcgctgctgatggtgatggccttggtcaccgcgctgtccacccaggtcaagggtacgctcacctacagccagttga
aagagcaggtgaagcaaggattggtgggtgacggcacgtcgccaaaagcgaaaactccacagtgaggattgttcaggg
caaggcgcagctttgttgagattatttttccggcatttttctacgatgactccaggcagtttgattgatgcggcctgcatgtcgc
gggctggttcgactgacgcttcaggagcattgccaagcgcaatccgtccaacatcatcgataccgcgtcgcgctggggca
tccgcagccgctcggcgctggtcaagggctaccgcaagcagttcaacgaagccccctccgaaaccatctggcgctgagc
cctccggcggaccgccgccgcgcccggtgcggcgcaggcagcggcggtccgctcccttcccatctcctggacaacatc
atgcatctgactcgagcgacgggccggaaggccggcgtgcggacttttcccgggcggctgaaaggcatggccgccgcg
ctgctgctatggacggcaggcacggtctgcgccgcgcccatcccctggcagtcgcagaaattcgaatacgtggccgacc
gcaaggacatcaaggaagtcctgcgcgacctgggggccagccatatgcaaaaaacccctcaagacccgtttagaggccc
caaggggttatgctagctgtcagatcgtcccattcgccatgccgaagcatgttgcccagccggcgccagcgaggaggctg
ggaccatgccggcctgagaccaatggtctcggttcacactccacaagctagctcgcaaaccgagcgcctgcccgaatac
ggttcacactccacaagctagctcgcaaaccgcgcgctgcagctggcgttgttcacactccacaagctagctcgcaaaccc
tgcgatacggtcggcgtcgttcacactccacaagctagctcgcaaacccttgtttatgagctatatgtgaagcgtccgtgcgg
tctttgtgcacggagtttcagccttttggctaccttaccgagcgttaccaaaccacttaacagttttattggcacttctcggcttat

cgtcgctgaaccgacgaatactagtattatacctaggactgagctagctgtcaatctagaccgccacggttgatgagagcttt
gttgtaggtggaccagttggtgattttgaacttttgctttgccacggaacggtctgcgttgtcgggaagatgcgtgatctgatc
cttcaactcagcaaaagttcgatttattcaacaaagccgccgtcccgtcaagtcagcgtaatgctctgccagtgttacaacca
attaaccaattctgattagaaaaactcatcgagcatcaaatgaaactgcaatttattcatatcaggattatcaataccatatttttg
aaaaagccgtttctgtaatgaaggagaaaactcaccgaggcagttccataggatggcaagatcctggtatcggtctgcgatt
ccgactcgtccaacatcaatacaacctattaatttcccctcgtcaaaaataaggttatcaagtgagaaatcaccatgagtgac
gactgaatccggtgagaatggcaaaagtttatgcatttctttccagacttgttcaacaggccagccattacgctcgtcatcaaa
atcactcgcatcaaccaaaccgttattcattcgtgattgcgcctgagcgagacgaaatacgcgatcgctgttaaaaggacaa
ttacaaacaggaatcgaatgcaaccggcgcaggaacactgccagcgcatcaacaatattttcacctgaatcaggatattctt
ctaatacctggaatgctgtttttccggggatcgcagtggtgagtaaccatgcatcatcaggagtacggataaaatgcttgatg
gtcggaagaggcataaattccgtcagccagtttagtctgaccatctcatctgtaacatcattggcaacgctacctttgccatgtt
tcagaaacaactctggcgcatcgggcttcccatacaatcgatagattgtcgcacctgattgcccgacattatcgcgagccca
tttatacccatataaatcagcatccatgttggaatttaatcgcggcctcgagcaagacgtttcccgttgaatatggctcataaca
ccccttgtattactgtttatgtaagcagacagttttattgttcatgatgatatatttttatcttgtgcaatgtaacatcagagattttga
gacacaattcatcgatgatggttgagatgtgtataagagacaggtccaaacaagcccgaaaacggcaccgtgctgatcga
cagcacagcagcagatctagcacctcccgttccgaaacgcctacagcaagcgctgcgtgcagtccaaacaagcccgaaa
acggcaccgtgctgatcgacagcacagcagcagatctagcacctcccgttccgaaacgcctgcatctagcaggcgttttttt
ttcgtgcggatctttcgatagtaccgatgccgagcggcatcctggtagccccggcgcagtgtccggatatgccagtacaga
tgcctgatctcgcgcgccacggccttcagttcgtcgtctgacactgccgattttggttgttctgagcagcgtagcatcggcac
gcgaagcgatccgccacagccagcgcaaccgaagatcgtaagcgtttcgaaagctggcttgcaggaaagggggtgtttg
tatgtgggcggccctgtatgtacgtacctggtcatacccgctgctcggtcccgccaggcgccctccgggcatcaaccgtcc
tgcacctggtcaactctcccatgtccccagaggggcccccttagcggctccgactacggcgcggacactcccgacgctgg
acgcaaaaaaaggccggcgatccgcacggcctttctatttttcgttaccttttcgttactccgggtagaactccggcgacagc
acctgctcgggtatccacggccactgctcggggaacacgtccaggccagtctcgtccacagccgccgagaccgcatccg
cccagatctcttccttccaatccggatcagccaacatcggctgcaggctgggcgtccgatgcaaccgcgccagaattgcgc
gacgctgttccttgatcgtgcgctgccagctcgaaccccggcggcccggctgatattgccacttgagcaaatgcgccagca
gtaccgccatgcggcttgccaactcccgctgttcgctcttgcccacgtcctcaatctcctccgcgatgtgccgaatgtcgatg
tctgaaagcttgccggcccgcaacagcgccgcctgctcactggcccacgccaccacgtccacttcgtagctcgttcccata
acgcgctccctccaataaccggatcattttagacgtttcgcgttacgccttgcgcgtaacgactttgcgatcaccataaattgc
ctcgccagccgcttgcatctgcgcctcactcacggccacagaacggacaaacgaagcgcgccacaactcggactcccgc
accttctggcgctgttcgaacgtattcgccttcgcagccaacgcgagcaagtccagcattgcgtcgaacgaaacgggatttc
cgttactcggcgtaacggatttttgctgcgactcaaacatcgcaaccttctcttccaagacgccaacataacgcaacgcttcct
gacgcccgtcctcgcacgtcataagctgccggcgcaacgcctcaatctccgcaagatttccgttactcagcgtaacggattt
ggcggcatcgctgatctcttccctcagccgcctattctcagcaaacgccttattgcgttcagcctccgccagctccacctggc
gacgcatgctcaccagatcagcgcgaaacttctcgacctccaggcggcattcatcgtaagcatccacgtcggaaagcattt
ccgttacggtcatacgcttagtaacggatttagcctcaacaggctgctgagccttacggcgagctcgatatgcggcttgacg
ctccgcgttggtcatagcgtgcgccttacgcggacgaccgcggccacgcctttgctccatccccggcagatcgaccgtca
cattatccgttacgtcacgcatattcggctcccgatttgatgattccattttacgttactggtaacggcaattcaattatcgttaca
atttagttcgtaacggaaatttcgcgagcagccccggcgctagacgccctcagcgggctttggagccgtttttctgactggg
cggtcgcaggggtcagctgcgccgaaattggctcctgcgccgcttgtggctcgcggtaggggtcaaacggcggcctcct
gatccactcccgacactgcatgtcatcgaggccggcatccgtgccttgagccgtgtagcacgtgcacctggtcgacgtgca
ggcaccgccgatcaccgtcggcatcgagcgaatctggcgcaactgcgcataggccggcgccgtctcaggccggccaga
gacagcagggacgaacgccgcgaggatctccgccccgtccgtcgccgcggcggacggacccgccttgaccgctaccg
ctagagctgaagccacaccgcccctgccctgctcggccgcaacaggcttagccggagtcgccgtcgtcaactggccggt

acggtaatagacccggtacgccaagaacgccgcgatcaacacgcaggcgatgaacagcatcagcaccggcggtacggt
gtacttgcgcttgatgtgcagactggaggacttgtacaggccgaaactggacttcggcaggctccacttcttcttgatcggcg
cggtgttgaacgtctccgggttcgcgcactccggccattcgtagtaccagcgtccgagcaagccagcgtcccgcaggtgg
acatgctgccccaccagcttgcggatatgactgtccaagaacgtcgggttctgcgtgatcagaacaaacgtcacgccggta
tgccgcaccgtctcaaacgccgccacgtggtcaggcaccttggacccggccgtgcggacgcgaaacacccgctgcgcc
tcatccaacacgatcagcgagttcggcgggaaggtgaagtacggcagcatcatccctgggttttcagggtcttcgcgcagc
tccgtccagtccgaaaccgccggcgtcgggatatacggcagcttcagctccgggatgcccatgacgaagagcggacgc
ccctggtcgacggctgccttcatcatctggaccgccaacgcggtcttcccgccaccaggcgtggccgtgatcagcgtgatc
ggttgcgttgcgctcatgtcagcttgcccagccgcttcagagtgatcatggagatgcgcgcggtgatgccgccggcgatga
tcgacagcccggtgaagacaccgccccgcgccaggatggccgcagcaacggcaggcatgccggccaggctactcttg
gccgcgcccagggcagcgctcaccgccgcatccaacccgacataggtgatcagaccgatcccgagcgacaccagtag
ctgacgcgcgagtggcccaacgagggccatgaggaacccagcgagcggcatcactcctcccctttacggcctacgccg
atgacgatcagcgcagcgcccaaccatgcgcacgcgatgatgaccggtcgaaacatgtcagcaccgtcacagaccggtt
tgagcgaccaagagatcggcatgccgtggatggaggcggtcagatcggacggacacggtgcggtatctgcgccccaac
cgctgtccggcatgaccttgacgttgacctgctgctccttcagatcgggcccgtccggtatctcgccttgctcgatacaaccc
atgcgcgtctcatggcccgagcactggtcgtcctgccgctccggagccttgccgttaccagtcgccggatccgtcgccgg
attgccgttcgcatccacctcctgcttagcggtcgtcaacgtggctgtcttgccatcggaatttggcgtgaccgtagcaacat
cgcggtaacgcttacctgtaacgggatcaacgtacgggtcgctcagattgacgttgactggagtggtggacggcgtgagct
tcaccggaatcggcaccttggctgcggccatatcgctagcaacagcagcgggcagcggatacgtcaaacctttgttccaat
ccgcatcgcttgcgcccaccgtcggaccggcagggtcgggcacacatgccgagccactcacgacatagccatcaacgc
agctggacgcctgactcgtgcccgcatagaaattgtcgccgccatcgttggtgtaatggcactcatagctcgtcccattacc
ggtagctttcatacccgcgaacttagccttctgcccagccaaatacgcgtctgccgccaacatcgcagagcacgcggcgc
caggagaagcagcaacgccattcgcgatattgccgccggtggcggacgtgttgtagccgtaattccagccccagccattg
aaccccgtgtcgccagcagccggtgaccgcttcgatgtgcaccacgtaccgtccaggcacttctggatgccgagctgcgc
caagtaggccaacgacgtcgcggtcgcgatcgcaggcgtcgcacgaagcgcggccagcgcaacggcagcagcgcctt
cgcccaccgtcatcgttgcggcgatcgtggcggcacctgcgccaaccgcggcacgcatctcggtagccgtcgcaacttc
agacagcgtcacagcaccagtcgtgacatacccagtccccgtcaacacgatgttcggcggcggaatcaacgggatggtc
gaagcccacgcggacgacgcccaccagcagagcagcagcacgagcacgcgcatcacagccccctgaaaatgattact
gccggcagaaccggcagcaggaatccggcccagagccagaaatcgatcgcgagcatcacagcccctttttcatcaccac
cagcgcccaggccgccaccatcgcggcaacgacgccccaccccatcgtcatgccatccttgaagctctcttgcgggtcac
atgccggaaacgacagcgacggcaacgacgcatcggtcaacgtgccgacgctcccatcactgcccacctgataccgcc
gcagcacccacccgcccgtggtcttgacgaactcagacagataggtgactgccccaggtgtctgcgacggagcgacgg
cgctgtaataggcatccgtggccatgcccacatcggcaaagcaccgtgcgcccaccaacgcgccgtcagcagccatcac
acgctccgacgcatgtacttgaacgtcgcaatgccgatcacgatgatcagcgccaggcccgccagcgtggcggtgtccg
ccttcgaatcggacatggcggtagcaacgtcggtcggcaccgcagccatcgccgaaccggccagtgcaacggtacccg
cggcaacagcagccgccttgctcttgatgctcttgaacatggttttctccttgagatggaggttgagaaaagctccgggccgt
tcaactcccagagccagggaacatcacgccttcttgggctcagcctcagcacgctgcagcggcttgatgctcgtaacgacc
ttctgaccgcccttatccttgccgttgctcgtctcgaccatcgagacttccgcgatgaacgggaacgggttgtggatgatcgc
cttcacaaccgcagagctctcgcacttcagctcctgcgtgcaggtgcccttcgagtcttcgccacgcagctccacatccgtg
tagatcttcccggtgtccagctgcttgccatccatgttgccaacccacgtcttggcgccccggatggtcacgcgtgcaatcat
ttccatggtttcactcctcaggtttcggcactggatcgtgcgtgccgtacacgtgcctcgccagggcggatttgtgcagctga
gccggtacgccctgacggcgtatcgcgacaaccagcgcagcgatgtcttcctctgtgcagcgcagctcgtaatcgaccgt
agggccgaactgcgtctggatgtgcttgagcttgcgttcgcggatggtctcgtcctgaagctccaaggccttgacctggtca
gtaggaacgcgctgcggatccgcagccatgaaggcctccagggccttgtacgcaccagcgaagtactggtcgcgcttga

taaggatttcgtgcggaatcacgcgatccttggcgccgaactcgatctcgagacgcacccactcgctatcctgattgccgag
ctggcggcccttctcgtaagcccgcagcatcttgccgttcgcccgacggccaatctcgaacgtcgtaccgcgacagccctt
gctgcccgccacgccgctctcgatcttgcgatacgtcgggatacgcccgcccgcgttgaagtcgccggcgtagtacagct
cttccatctgcgcgatgctcacctcgccctggcagaagtccatcgccaggtcgcaccgcgtgatccgcgcgtcgaggtcct
gcaccatcgcgtagacggcttgccagtcgccaatcgcggtgcagccctgccccggccagtccaccaggatcgtgccgcc
aacgtgctcgccgccgcaggcaacgatcccgagcttcatcgtctcgccgttgatgaacgccagcaggtcgtagctgaact
cataccggcggaaccctttgcccgcaggcttcatcgtcaccggcaccgagaacaccagctggaagtacctgcgcagctg
ctccagggcgtcgctgatgctgccgtcgggcaggaacgtgaacttgaaccagtccacgatcgcacctgccttgcgttctgg
actttccccgggttttaccggacccggggaacggcctgtcggccgcccagcctcgctccgctcggcccggtcggccgcc
atgccgttctcgcgtgcgagcgccggcccatgaactccatacgcaatgcctgtgggatcaagatgacgagagggggtgg
gacggacgcgcatcagcgtgcgccctcgcagcgataccagttcgccagatcggcctggcgcgagttcttcaggaggtcg
tcaatgcgatgcatcgcaggacgcgattgcaggcagcaggcgccgcgcagaacggcagccatcacgatggccagctgc
gcaccgtgcgggtgaagcacgtcgccaatcaggtccagctgcctatggtctctctgctgcat
